# Principles of Computation by Competitive Protein Dimerization Networks

**DOI:** 10.1101/2023.10.30.564854

**Authors:** Jacob Parres-Gold, Matthew Levine, Benjamin Emert, Andrew Stuart, Michael B. Elowitz

## Abstract

Many biological signaling pathways employ proteins that competitively dimerize in diverse combinations. These dimerization networks can perform biochemical computations, in which the concentrations of monomers (inputs) determine the concentrations of dimers (outputs). Despite their prevalence, little is known about the range of input-output computations that dimerization networks can perform (their “expressivity”) and how it depends on network size and connectivity. Using a systematic computational approach, we demonstrate that even small dimerization networks (3-6 monomers) are *expressive*, performing diverse multi-input computations. Further, dimerization networks are *versatile*, performing different computations when their protein components are expressed at different levels, such as in different cell types. Remarkably, individual networks with random interaction affinities, when large enough (≥8 proteins), can perform nearly all (∼90%) potential one-input network computations merely by tuning their monomer expression levels. Thus, even the simple process of competitive dimerization provides a powerful architecture for multi-input, cell-type-specific signal processing.

**Graphical Abstract:** 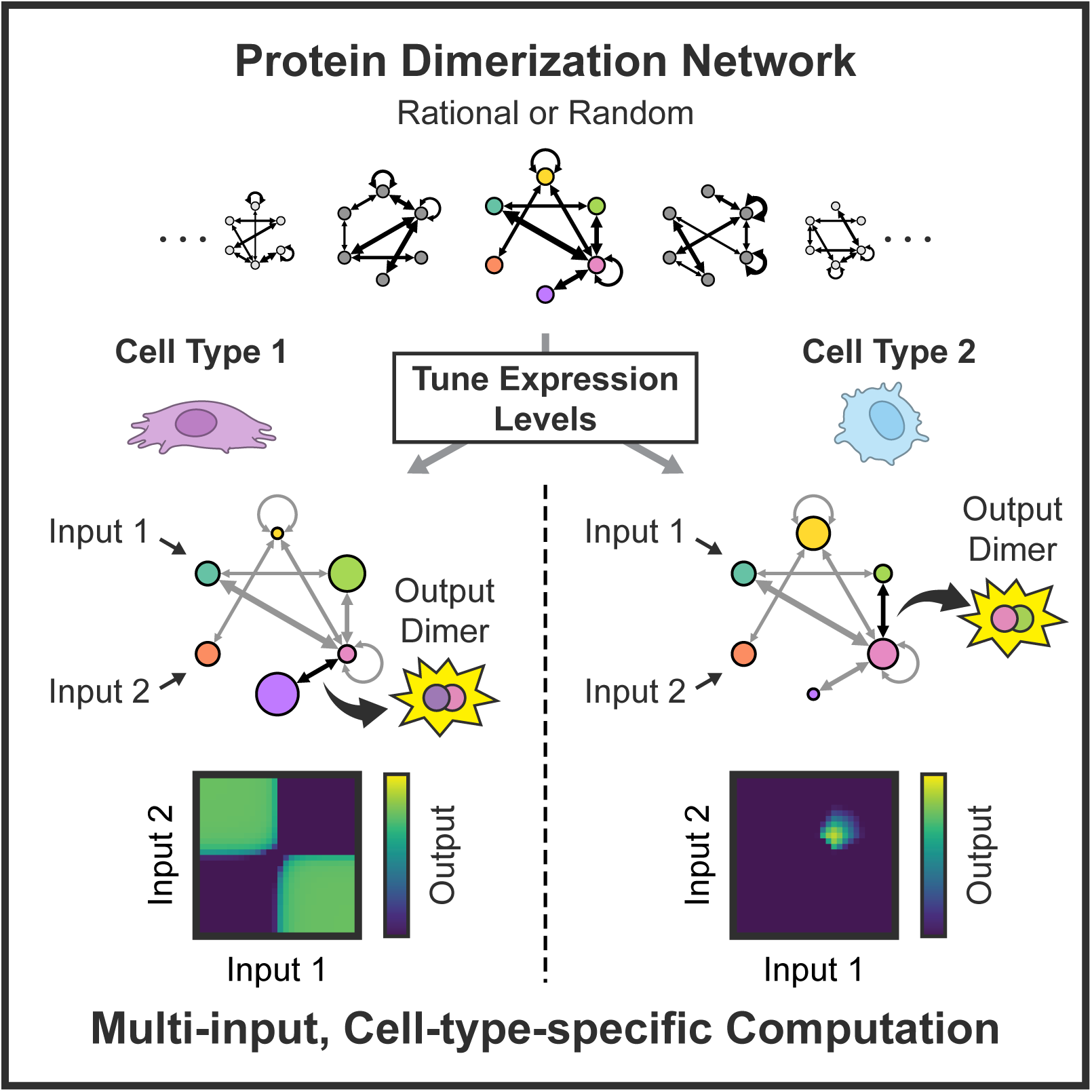

## Introduction

Many biochemical signal processing circuits employ families of proteins that competitively dimerize with one another in diverse combinations. For example, the motif of many-to-many dimerization can be found in transcription factor families such as the nuclear receptor (NR)^1^ (Figure 1A), basic leucine zipper (bZIP),^2–4^ basic helix loop helix (bHLH),^5^ and MADS-box proteins,^6^ among others.^7^ These dimerization networks integrate a variety of signals from other cells, environmental cues, and the intracellular state to regulate genes involved in major cellular decisions such as cell proliferation,^8–10^ differentiation,^11–13^ and stress responses.^14^ Similar dimerization (or higher-order multimerization) networks also occur in ligand-receptor signaling,^15–17^ adhesion,^18,19^ and other systems.^20^ In addition to their many-to-many patterns of dimerization (Figure 1B), these proteins commonly exhibit diverse but overlapping expression profiles across different cell types,^21^ calling into question how these cell types might interpret input signals differently (Figure 1C).

**Figure 1.**
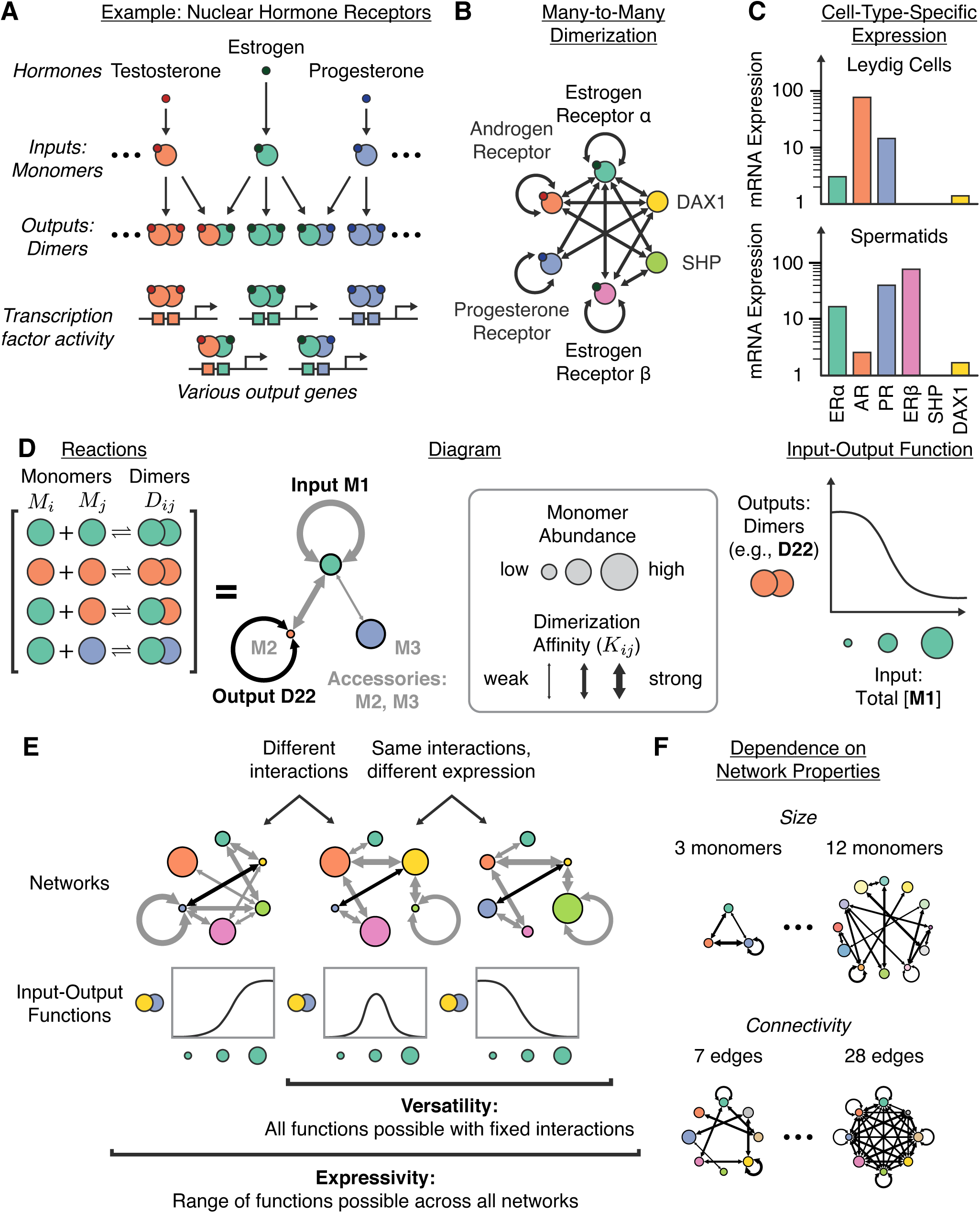
Competitive dimerization networks compute. (A) In natural dimerization networks, such as the nuclear receptors (NRs), upstream signals regulate the activities of individual monomers, which dimerize to perform biochemical outputs. (B) Network monomers dimerize in a many-to-many fashion, as shown for several NRs.^1^ (C) Different cell types, such as spermatids and Leydig cells, express NR monomers at different abundances, reported in normalized transcripts per million (TPM) from the Human Protein Atlas.^125^ (D) Dimerization networks are modeled here with pairwise affinities (shown as arrow widths) and monomer expression levels (shown as circle sizes) as parameters. Titrating input monomer(s), with a dimer as an output, yields an input-output function. (E) Here, expressivity refers to the collection of input-output functions that may be performed by all networks of a given class, while versatility refers to the functions that may be performed by a single set of proteins with fixed interactions but variable expression levels. (F) This work investigates how network expressivity and versatility scale with both network size and connectivity.

The process of competitive dimerization can be thought of as a biochemical input-output computation.^22^ In this perspective, upstream signals modulate the concentrations or activities of specific monomers (inputs), altering the distribution of biologically active dimers (outputs).^23,24^ For example, steroid hormones bind and activate cognate nuclear receptors (NRs), allowing them to dimerize with other NR monomers (Figure 1A). The resulting dimers can bind to sites across the genome to regulate downstream genes. In this work, we use the terms “computation” and “response function” to refer to the quantitative relationship between the concentrations of input monomer(s) and an output dimer.

In 1993, Neuhold and Wold suggested that networks of dimerizing bHLH transcription factors could allow changes in just a few monomers (inputs) to “radiate” throughout the network, changing the concentrations of dimers (outputs) to generate “major changes in cellular phenotype.”^25^ Since then, signal integration and decision-making by dimerization networks have been documented in many other contexts. In lymphocyte development, two sets of bHLH proteins – E protein transcription factors (E12, E47, E2-22) and inhibitory Id proteins (Id2, Id3) – dimerize with one another to control multiple decisions, including the choice between innate and adaptive immune cell fates.^11,26,27^ These bHLH proteins are regulated by pre-T-cell receptor (pre-TCR) signaling,^28,29^ Notch signaling,^30,31^ and certain cytokines.^32^ Further, the effects of E proteins on gene expression are contextual, varying across developmental stages.^11,27^ Another example occurs in the BCL-2 family of apoptotic proteins, where BAX and BAK proteins homo-oligomerize to form pores in the mitochondrial outer membrane, inducing apoptosis.^33^ Anti-apoptotic proteins in this family (e.g., BCL-2, BCL-X_L_) heterodimerize with BAX and BAK, preventing their homo-oligomerization, whereas pro-apoptotic BH3-only proteins (e.g., BAD, BID) can bind to and inactivate the anti-apoptotic proteins. Cellular stresses, such as DNA damage,^34,35^ hypoxia, and oxidative stress,^36^ as well as survival signals – such as those used in lymphocyte development^37–40^ – regulate the balance between the pro-and anti-apoptotic proteins to control pore formation and apoptosis.^41^ Finally, dimerizing *Arabidopsis* bZIP transcription factors in the C and S1 families^42–45^ integrate signals from “low-energy” abiotic stresses (such as drought, darkness, salinity, and hypoxia, which all reduce sugar abundance) to slow growth and induce changes in metabolism.^44,46–50^ For instance, sucrose translationally represses all S1-family bZIPs,^51,52^ glucose represses the transcription of bZIP1 and bZIP63 through multiple mechanisms,^47,53^ and the stress response kinase SnRK1 activates bZIP63.^54^ Different tissues, such as roots and leaves, express the bZIP proteins at different levels,^43,55^ allowing them to play distinct roles in the plant’s metabolic response to stress.^49,56–59^ Thus, dimerization networks appear in diverse biological systems, integrate multiple inputs, and operate contextually.

Despite their prevalence and significance, the computational capabilities of dimerization networks remain poorly understood. Dimerization is a relatively limited type of biochemical interaction that does not consume energy and is stoichiometric rather than catalytic. In contrast to enzymatic networks^60–63^ and transcriptional regulation,^64,65^ dimerization is incapable of amplifying the magnitude of input signals.^23^ While the experimental characterization of particular dimerization networks in nature has provided great insights,^11^ and some dimerization networks have recently been studied computationally,^66–68^ we lack a fundamental, systems-level understanding of dimerization network computation – including which computations are (and are not) possible, to what extent a single network can perform different computations in different cell contexts, and how parameters such as network size and connectivity influence their computational power. Addressing these questions is essential for understanding the prevalence, architectures, expression patterns, and signal-processing functions of natural dimerization networks, as well as for engineering synthetic dimerization networks.

Here, to study dimerization network computation in general, we constructed a minimal model that captures the key features of natural dimerization networks: competitive, many-to-many dimerization interactions of varying strengths and cell-type-specific component expression levels (Figure 1D). We adopt the term *expressivity* from the field of neural network computation to describe the range of all quantitatively unique functions that may be performed by a class of dimerization networks across all physiologically reasonable parameter values (Figure 1E).^69,70^ Further, we use the term *versatility* to describe the ability of the same network of proteins to perform different functions when network proteins are expressed at different abundances (such as in different cell types). Network versatility would allow different cell types to reuse the same set of proteins to perform different modes of signal interpretation.

In this work, we use both random parameter screens and optimization to characterize the expressivity and versatility of competitive dimerization networks. We find that dimerization networks can compute a variety of non-monotonic functions on multiple inputs. We investigate how network expressivity and versatility vary with network size and connectivity (Figure 1F) and use our results to contextualize the features of natural dimerization networks. We then demonstrate that dimerization networks can readily perform computations on multiple inputs, including all three-input logic gates. Finally, we show that even networks with random protein-protein interaction affinities, when large enough, can perform a wide variety of functions solely by adjusting the expression levels of their monomer components.

## Results

### Modeling competitive dimerization networks

We first sought to establish a minimal modeling framework that captures the key features of natural dimerization networks described above: competitive, many-to-many dimerization interactions of varying strengths and variation among cell types in component expression levels. We consider networks of *m* interacting monomers M_1_, M_2_, … M*_m_*. Each pair of monomers M_i_ and M_j_ may reversibly bind, with equilibrium constant K_ij_, to form the dimer D_ij_ (Figure 1D).

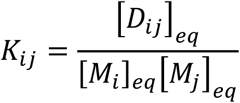

The total concentration of each species is the sum of its free form and all of its dimers:

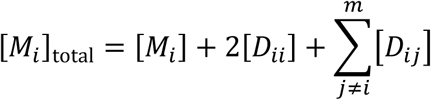

To consider these networks as feed-forward computational systems, we designate a subset of monomers as *inputs.* The expression level of an input monomer depends on a corresponding input signal. We term non-input monomers *accessories* and assume that each accessory protein has a fixed total concentration in a given cell type. We consider dimer concentrations as the outputs directly, assuming that cells could use various dimerization-dependent biochemical activities to carry out downstream functions.

In this framework, a given network in a given cell type is completely specified by its set of pairwise affinities, K_ij_, and the total concentrations of each of the accessory monomers, [M_i_]_total_. To simulate the input-output function of a network, we determine the equilibrium concentrations of all network species across a titration of the input monomer(s) (see Methods). This framework does not consider the formation of higher-order oligomers, such as trimers,^71,72^ and also neglects the potential impacts of DNA binding or subcellular localization on dimerization propensities.^7^ These features could further expand the computational potential of these systems beyond what is described below.

### Networks of dimerizing proteins can compute a wide variety of functions

What types of functions can dimerization networks compute? To address this question, we first analyzed the input-output behaviors of minimal, elementary networks. The simplest non-trivial dimerization network comprises just two monomers (Figure 2A). In this network, increasing the total concentration of M_1_ induces the formation of D_12_ heterodimers, which in turn sequesters M_2_ and prevents the formation of the D_22_ homodimer. Thus, this network computes a *switch-off* function with output dimer D_22_.

**Figure 2.**
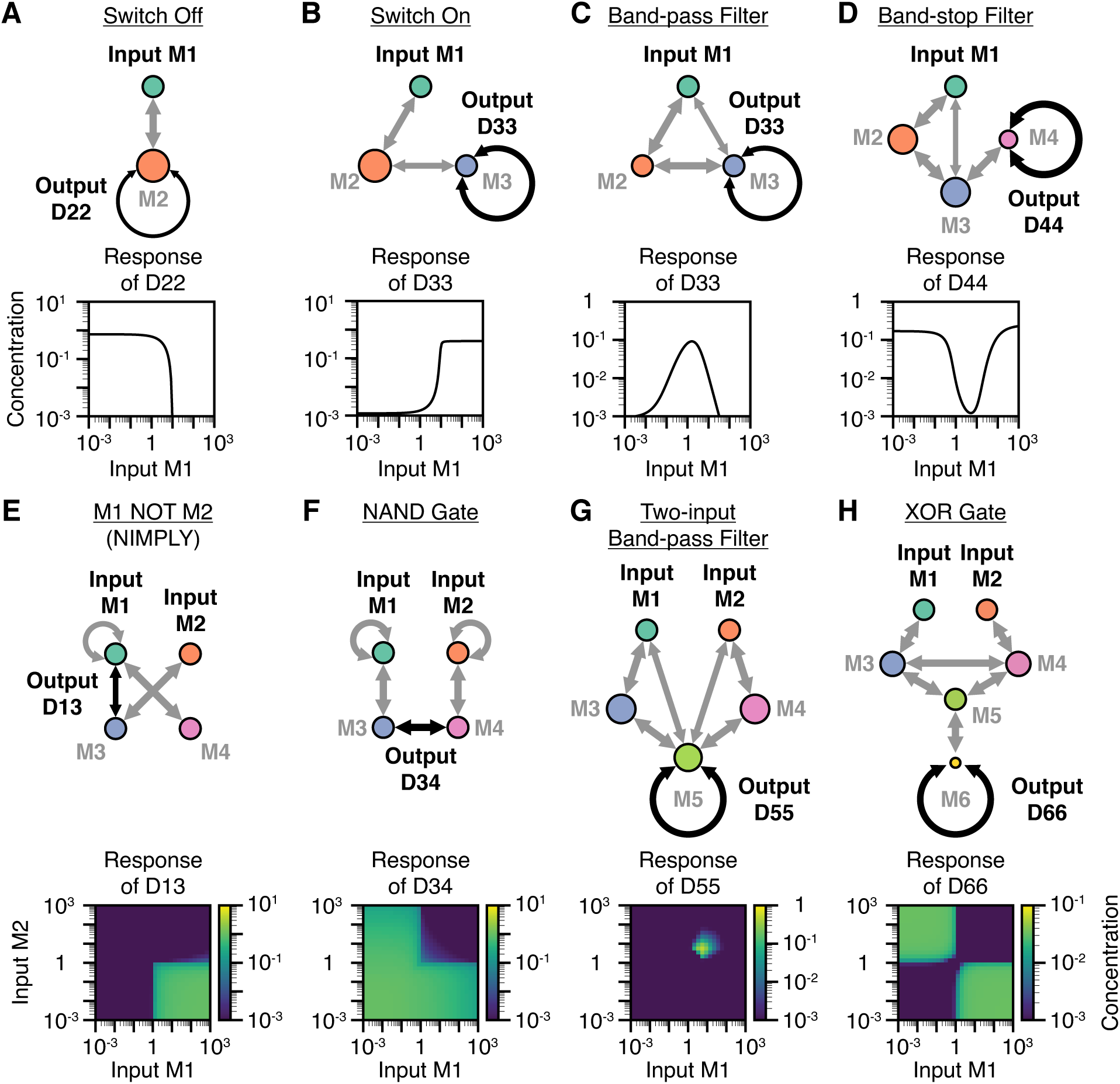
Competitive dimerization networks can compute diverse functions on one and two inputs. (A-D) Examples of one-input, one-output functions. From left to right, simulations of networks performing a switch-off function (A), a switch-on function (B), a bump function (C), and an inverted bump function (D) are shown. (E-H) Examples of two-input, one-output functions. From left to right, simulations of networks performing an M1 NOT M2 (NIMPLY) gate (E), an M1 NAND M2 gate (F), a two-input bump function (G), and an M1 XOR M2 gate (H) are shown. For all panels, the networks shown were inspired by networks from the random parameter screen (Figure 4) and rationally pruned to identify minimal topologies capable of computing each input-output function. All input-output functions are displayed in unitless concentrations (see Methods).

Adding an additional species can invert this circuit to compute a *switch-on* function (Figure 2B). In such a network, increasing the input monomer M_1_ increases D_12_ dimers, reducing D_23_ dimers, ultimately increasing the D_33_ dimer. This example illustrates the way in which concentration changes can propagate through a network.^23,73^

When the switch-on and switch-off networks are combined, a biphasic *bump* function emerges, in which only intermediate concentrations of the input monomer promote formation of the output dimer (Figure 2C). In this network, the input M_1_ dimerizes strongly with M_2_ and weakly with M_3_. M_2_ and M_3_ strongly heterodimerize, and M_3_ homodimerizes to form the output dimer, D_33_. As total M_1_ levels increase, they initially form D_12_ heterodimers, thereby decreasing D_23_ heterodimers and allowing the formation of the D_33_ output dimer. However, as M_1_ increases further it begins to form dimer D_13_ as well, thereby suppressing the formation of the D_33_ output dimer. Thus, the output dimer D_33_ can form only when M_1_ is present at intermediate concentrations. As with the switch-on and switch-off functions, adding an additional monomer can invert the response to produce an *inverted bump* function that responds only outside a window of intermediate input concentrations (Figure 2D). Taken together, these results provide an intuitive picture of how dimerization networks of increasing size can compute functions of increasing complexity.

Many natural signaling networks respond to multiple input signals,^15,74^ provoking the question of whether dimerization networks can similarly compute functions on multiple inputs. Indeed, by considering two monomers as distinct inputs, combinatorial logic can emerge. For example, Figure 2E shows a network that computes a NIMPLY (“M_1_ AND NOT M_2_”) logic gate. Here, the output dimer D_13_ forms in the presence of input M_1_ alone, but not in the presence of input M_2_, which competes to dimerize with monomer M_3_. Similarly, Figure 2F shows a network computing a NAND logic gate, in which the output dimer D_34_ forms in the absence of inputs or the presence of a single input, but not in the combined presence of both inputs M_1_ and M_2_. Dimerization networks can also generate analog (non-Boolean) combinatorial responses. For example, the five-monomer network shown in Figure 2G combines two bump function motifs (Figure 2C) to create a two-input bump function, in which the output dimer D_55_ only forms in the presence of intermediate amounts of both inputs. Finally, larger networks can perform functions with even greater complexity. For example, a six-monomer network can compute an XOR logic gate (Figure 2H). Other functions beyond those described here are possible and can be rationally understood; atlases of networks performing one- and two-input computations can be found in Figure S1 and Figure S2.

This exploration of elementary dimerization networks demonstrates three key features of computation by dimerization. First, even relatively small networks can perform non-monotonic computations on one or multiple inputs. Second, many dimerization networks can be intuitively understood by analyzing the paths by which input perturbations propagate to affect output dimers. Third, networks of increasing size appear capable of performing increasingly complex input-output responses.

### Monomer expression levels can modulate network computations

A key feature of competitive dimerization networks is their computational versatility, defined as the ability of a single network to perform distinct input-output computations depending on the expression levels of its protein components. For example, the bump function shown previously in Figure 2C can be tuned by modulating the expression levels of the accessory monomers M_2_ and M_3_ (Figure 3A). In this case, the total abundance of M_2_ tunes the center of the bump because the input M_1_ must sequester M_2_ to induce the formation of the output dimer D_33_. Additionally, the abundance of M_3_ tunes the amplitude of the bump by directly promoting D_33_ formation. In another three-component network, adjusting protein expression levels can change a switch-off to a switch-on response (Figure 3B). Thus, even simple networks can show both quantitative and qualitative versatility.

**Figure 3.**
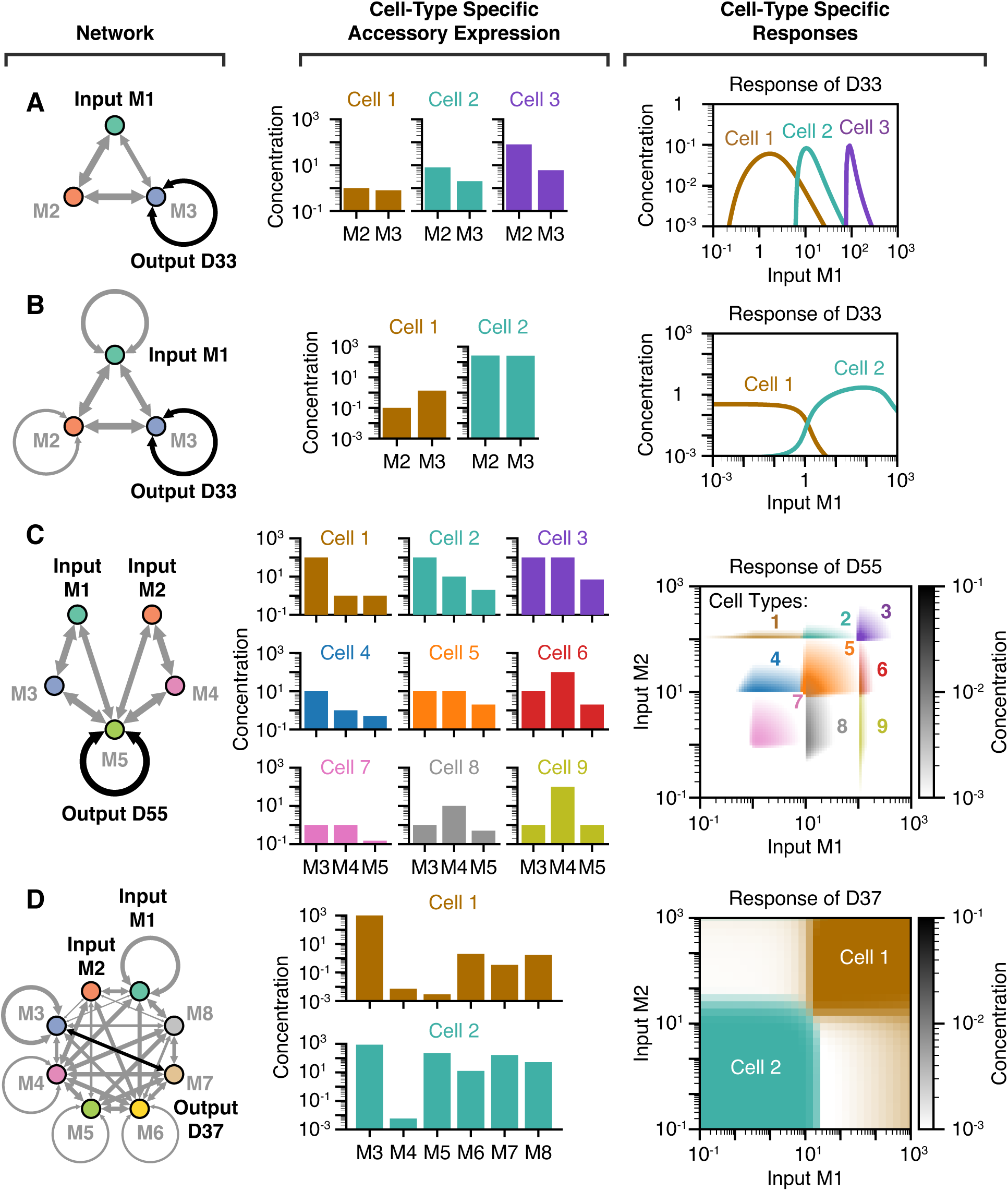
A single set of dimerizing proteins (left), when expressed at different abundances (middle), can perform different input-output functions (right). Accessory monomer expression levels can be used to (A) tune the midpoint of a bump function, to (B) transform a switch-on function to a switch-off function, to (C) tune the midpoint of a two-input bump function in both input dimensions, or to (D) transform an AND gate into a NOR gate. The network shown in (D) was identified by screening random interaction affinities for networks of eight monomers (see “Large random networks are expressive and versatile”). All input-output functions are displayed in unitless concentrations (see Methods).

Computational versatility extends to multi-input functions. For example, the midpoint of the two-input bump function shown previously in Figure 2G can be tuned by independently adjusting the concentration thresholds for inputs M_1_ and M_2_ (Figure 3B and Supplementary Video 1). Biologically, these functions would allow different cell types to respond to different combinations of two inputs, a concept known as addressing.^75^ A single network can even be tuned to perform entirely different types of multi-input combinatorial logic. For example, the network shown in Figure 3D and Supplementary Video 2 computes an AND gate with dimer D_37_ with one set of monomer expression levels, but a NOR gate with another. A different network can compute both OR and NOR logic gates (Supplementary Video 3). These examples of two-input versatility were identified simply by screening a small number of networks with random affinities, as described in the subsequent sections. Thus, these examples represent just a fraction of the potential versatility of dimerization networks.

### Expressivity grows with network size and connectivity

Evidently, dimerization networks can perform complex and versatile computations. How do the computational capabilities of dimerization networks scale with their size and connectivity? To address this question, we combined computational screening with optimization trials to systematically analyze the scope of possible network computations. First, we simulated large sets of networks of different sizes, with randomly chosen affinities and component expression levels. To generate each network, we first randomly generated connected graphs with varying numbers of edges (heterodimerization interactions) and then sampled the particular affinity values for each edge. We sampled parameter values from broad ranges consistent with experimentally measured affinities and protein expression levels (see Methods). Overall, this screen included 1 million networks of each network size, from *m*=2 to *m*=12 monomers.

For each random network, we determined the equilibrium concentrations of all species across a titration of the input monomer(s). By considering each dimer as a possible output, each network simulation produced multiple input-output responses. In order to classify these responses, we introduced a gridding scheme, binning the input and output into segments of equal size in logarithmic units (Figure 4A). With this scheme, two responses that pass through the same boxes are classified as the same function type. This allows a more tractable, discrete analysis of the continuous input-output response space.

**Figure 4.**
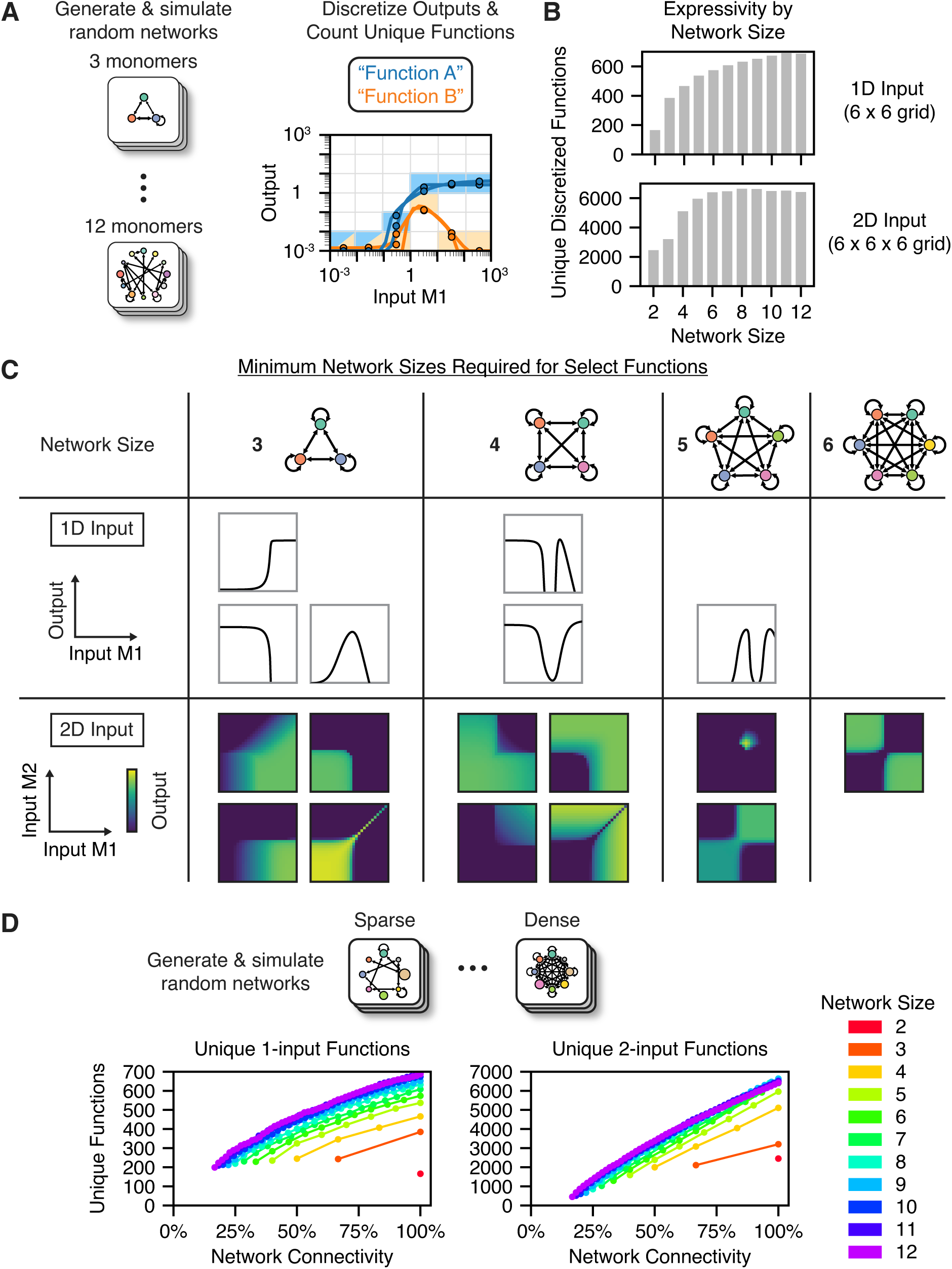
Dimerization network expressivity grows with both network size and connectivity. (A) *n*=10^6^ networks of each network size with randomized parameters were simulated and their input-output responses were categorized by discrete binning. (B) A bar graph shows how computational expressivity, measured as the number of unique, discretized response functions observed, increases with network size for both one-input and two-input functions. (C) Shown are the classes of response functions that become possible as network size is increased from three to six monomers, as determined by parameter optimization. See Figure S1 and Figure S2 for schematics and quantitative plots of each response function shown. (D) A plot shows, for different network sizes, how network expressivity grows with network connectivity, defined as the fraction of possible heterodimerization interactions in a network.

Unsurprisingly, larger networks performed more unique one-input and two-input functions (Figure 4B). For one-input functions, larger networks also exhibited a larger fraction of non-monotonic responses, including functions with up to 4 local extrema (Figure S3A). However, beyond network sizes of about *m=*6, we observed diminishing returns in the number of unique functions discovered, and this trend was not impacted by the number of networks simulated (Figure S3C, Figure S3D). In larger networks, fewer dimers successfully form and respond to input perturbations (Figure S3B), reflecting quadratic growth in the number of potential dimers, which increases competition for the limited number of monomer constituents. Additionally, for a given network size, networks with higher connectivity (density of heterodimerization interactions) produced more unique functions in the parameter screen (Figure 4D).

Within a network, each dimer represents a potential output. Can a single network use different dimers to compute multiple output functions? Or, does indirect coupling among dimers limit the repertoire of functions a single network can compute? To assess the range of possible two-output functions, we counted the number of unique two-output functions observed for every combination of dimers in every network of the parameter screen (Figure S3G). We then compared this number to a scrambled control in which an equal number of responses from the overall dataset were randomly paired together. Strikingly, the number of observed two-input functions closely approached that of the scrambled control, increasing with network size for both one-input (Figure S3H) and two-input functions (Figure S3I). This suggests that most combinations of two response functions can be implemented by two dimers in the same network.

While screening many random networks allows for an unbiased exploration of possible response functions, this approach could fail to discover certain functions because of finite sampling depth. Thus, to gain more insight into the network size requirements for specific functions of interest, we turned to optimization approaches to identify parameter values that compute desired target functions. We found that both simulated annealing^76^ and genetic algorithm^77^ optimization could successfully optimize network parameters (see Methods). By defining an error threshold constituting a “successful” optimization, we optimized networks of decreasing size until no satisfactory parameter set could be identified. For example, while optimization could consistently identify four-monomer networks performing an inverted bump function, they consistently failed to optimize three-monomer networks, suggesting that the inverted bump function requires four network monomers. This approach revealed the network size requirements for the one- and two-input functions shown in Figure 4C, Figure S1, and Figure S2. Together, the results of the parameter screen and optimizations suggest that most one- and two-input dimerization network computations can be performed by networks of just six monomers and, more generally, shows how expressivity grows as more monomers are added to a network.

### Competitive dimerization networks can compute multi-input functions

Cells commonly respond to many signals from other cells, their environment, and their own cell state.^15,74^ For example, cell fate decisions can depend on inputs from multiple developmental signaling pathways, which may activate a common set of dimerizing transcription factors.^11,32^ To understand how competitive dimerization networks could compute responses to multi-input signals, we used optimization to identify networks that perform various three- and four-input Boolean functions (logic gates).

Dimerization networks were capable of computing diverse multi-input functions. For example, a network of ten monomers can compute an “any 2 or none” function, in which the output dimer is formed if exactly two (but any two) inputs are present or no inputs are present (Figure 5A). This function would be difficult to implement by connecting elementary two-input functions, as it would require at least 11 AND and OR gates functioning orthogonally (Figure 5B). Thus, dimerization networks appear to offer the ability to compute multi-input functions in one computational layer, without the need for many orthogonal components. Using optimization trials, we demonstrated that dimerization networks of ten monomers can perform all 54 unique three-input logic gates, and networks of just six monomers could compute half of such functions (Figure 5C, Figure S4A).

**Figure 5.**
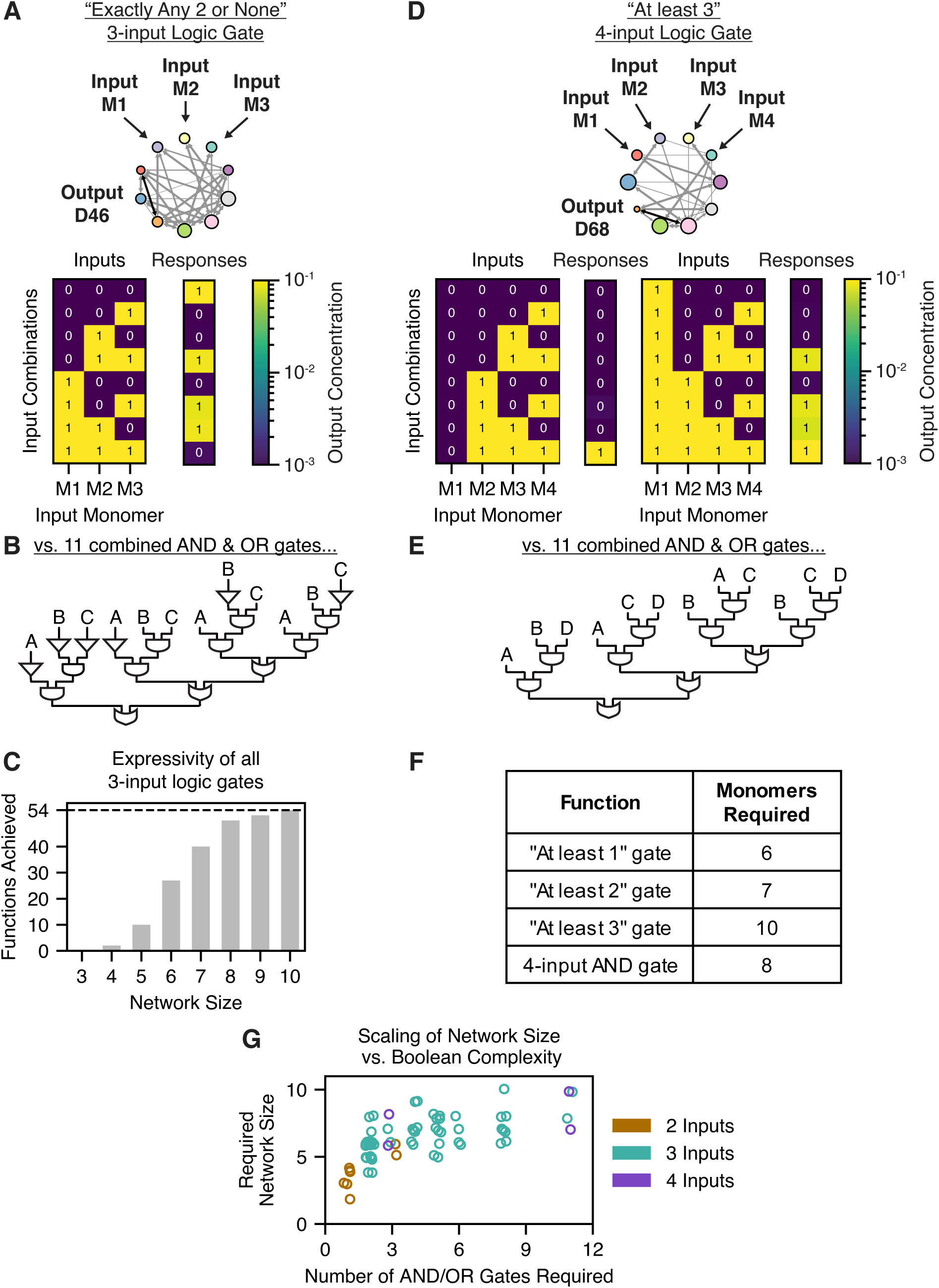
Competitive dimerization can integrate multiple inputs in multi-dimensional response functions. (A) A schematic is shown of a network computing a three-input logic gate, the “exactly any 2 or none” logic gate (explained in the text). Below is a heatmap of the simulated output concentrations for various input combinations (i.e., the truth table). (B) A schematic is shown decomposing the “exactly any 2 or none” logic gate into 11 elementary Boolean AND and OR gates. (C) A bar graph shows how the number of achievable three-input logic gates grows with network size. (D) A schematic is shown of a network computing a four-input logic gate, the “at least 3” logic gate (explained in the text). Below is a heatmap of the simulated output concentrations for various input combinations. (E) A schematic is shown decomposing the “at least 3” logic gate into 11 elementary Boolean AND and OR gates. (F) A table displays the number of monomers required to compute the four-input Boolean “at least” functions. (G) Shown is a graph showing how the required network size grows with the Boolean complexity (measured as the number of elementary AND and OR gates required) of various logic gates. Colors denote the number of inputs used in each function. A jitter (random perturbation) was applied to each point to distinguish overlapping points.

This result encouraged us to ask whether dimerization networks could perform Boolean functions on even more inputs. We thus defined a set of four-input Boolean functions, which we call the “at least *n*” functions, whose output is high when at least *n* inputs, but any *n* inputs, are present. For instance, the network shown in Figure 5D computes an “at least 3” function using ten monomers. This function would also be difficult to achieve using elementary two-input functions, requiring 11 AND and OR gates at minimum (Figure 5E). We found that dimerization networks could readily perform all of the four-input “at least *n*” gates as well as a four-input AND gate (Figure 5F, Figure S4B).

Thus, competitive dimerization networks can integrate several inputs in multi-dimensional computations. Importantly, while such computations could be performed using many orthogonal two-input logic operations, dimerization networks perform them all in the same computational layer. Indeed, while increasingly complex Boolean functions require larger dimerization networks – defining complexity by the number of two-input AND and OR gates they require – this relationship appears nonlinear, eventually reaching a point at which fewer monomers than elementary logic gates are required (Figure 5F).

### Networks adapt over timescales of minutes to hours and function in the presence of noise

Competitive dimerization networks appear to compute diverse functions based on the equilibrium modeling described above. But how rapidly do these networks approach steady state, and can they continue to function despite various sources of biological noise? Deterministic simulations with physiologically reasonable parameters (Methods) suggest that networks re-equilibrate on a timescale of 100 s to 2 h following a perturbation of the input monomer (Figure S5A), with only modest dependence on network size. This timescale, which is comparable to the dissociation timescales of the highest-affinity dimers (*k*_off_ ≈ 10^-4^ s^-1^ gives *t*_1/2_ ≈ 1.9 h) is faster than the hours to days timescales associated with transcriptional regulation.^78,79^ To test whether networks could remain at quasi-equilibrium as inputs change over time, we simulated network dynamics as the total concentration of one monomer was oscillated at varying frequencies. When the input monomer oscillated with a period of 27 h, over 80% of dimers remained near equilibrium (within 3-fold), whereas only 20-60% of dimers were at equilibrium when the same monomer oscillated every 100 s (Figure S5B). These results suggest that dimerization networks can remain at quasi-equilibrium when their components change on physiologically relevant timescales of hours to days.

Biological circuits must be robust to various sources of both intrinsic and extrinsic noise.^80,81^ In a cell, a competitive dimerization network would face both intrinsic noise from stochasticity in the dimerization equilibrium itself and extrinsic noise from fluctuations in the expression levels of network proteins. To characterize the impact of intrinsic noise, we performed stochastic Gillespie simulations of dimerization networks at equilibrium. Most species exhibited a noise coefficient of variation less than that of protein expression levels (0.2 to 0.5)^80,82^ (Figure S5C, Figure S5D). Low-abundance dimers were the most sensitive to intrinsic noise (Figure S5D), consistent with previous work,^83,84^ whereas most species with abundances above 100 molecules per cell exhibited noise coefficients of less than 0.1 (Figure S5C). To measure the effects of extrinsic fluctuations in monomer expression levels, we simulated networks performing each unique response function from the random parameter screen with 50 random perturbations of the accessory expression levels (see Methods).^80,82^ Most functions were robust to such perturbations, with the median root-mean-square deviation (RMSD) in the log-scaled input-output function being 0.2 to 0.4, corresponding to a 1.5-fold to 2.5-fold change in the output (Figure S5E, Figure S5F).

### Large random networks are expressive and versatile

It is unlikely that all the pairwise interaction affinities of a network can be precisely fine-tuned over evolution through protein mutations. However, protein expression levels could be tuned independently through several mechanisms, such as by adjusting promoter strength.^85,86^ In the field of neural computation, it has been shown that sufficiently complex networks with random weights can still achieve arbitrary levels of expressivity, provided that only the weights of the final output layer are tuned.^87–90^ Could dimerization networks with random interaction affinities similarly perform a wide variety of functions if only the expression levels of their accessory monomers can be tuned?

To address this question, we asked whether random networks could compute sets of various one-input target functions (Figure 6A). We generated 50 networks with random interaction affinities at each network size from *m*=2 to *m*=12 monomers. As target functions, we selected representative subsets of the one-input functions previously identified in the parameter screen for networks of each size. We systematically optimized the accessory monomer expression levels for each possible output dimer in each random network to best fit each of the target functions. We used the Chebyshev distance to determine whether a particular target function was “achieved,” requiring that the optimized response be within 10-fold of the desired target function at every concentration of the input monomer (see Methods).

**Figure 6.**
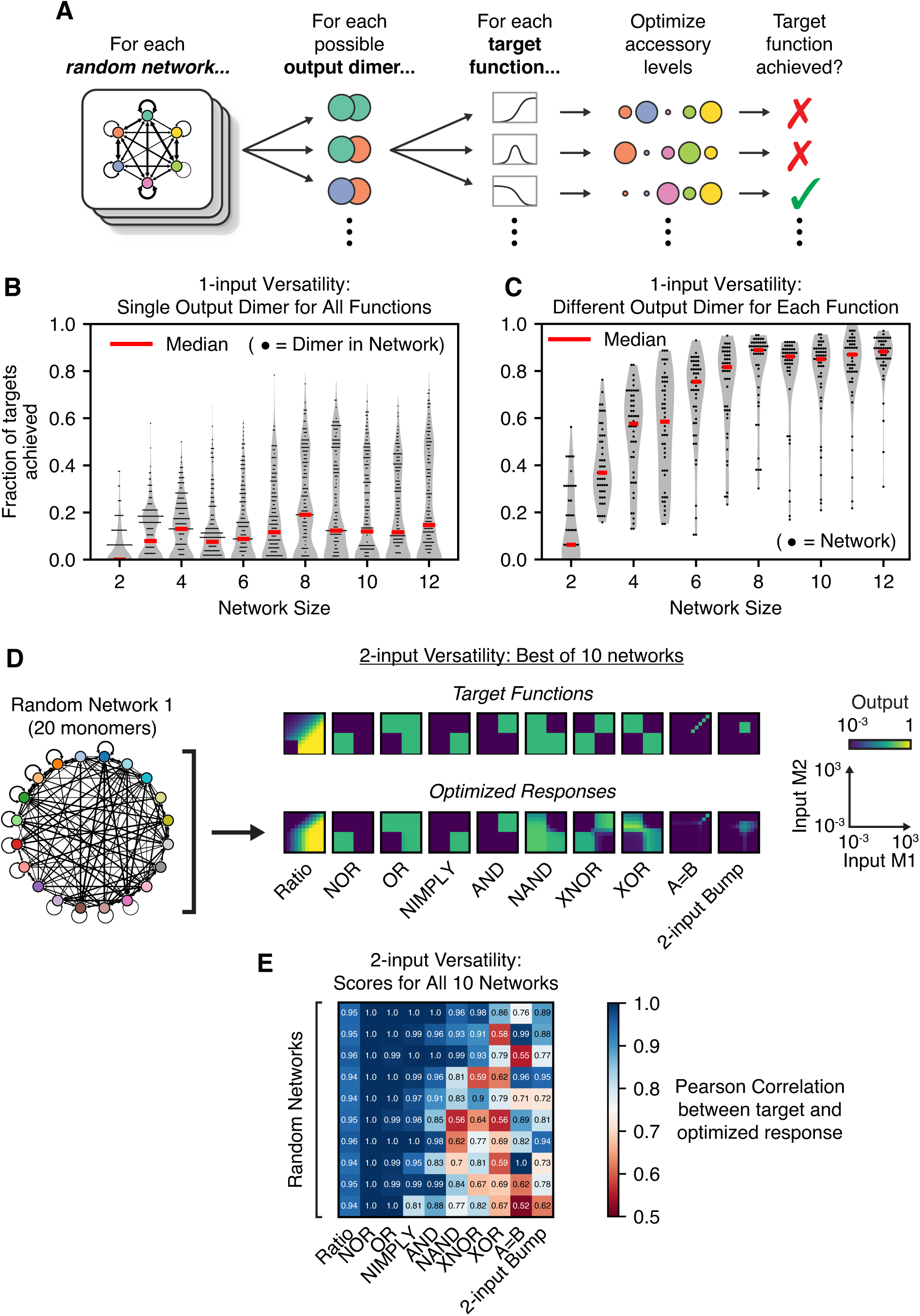
Large random networks are expressive and versatile. (A) For each possible output dimer in each network with random interactions (*n*=50), accessory monomer expression levels were systematically optimized to best fit each of the target functions. (B) A violin plot with scattered points shows the versatility of individual dimers, measured as the fraction of targets each individual dimer in each random network could achieve, for each network size tested. (C) A violin plot with scattered points shows the versatility of random networks, for each network size tested, when different output dimers may be used for each target function. (D) An example of 2-input versatility, showing the most versatile of ten tested random networks. Shown is both a schematic of the network (left) as well as heatmaps of the simulated responses (bottom row) after its accessory expression levels were optimized to perform each target function (top row). (E) A heatmap shows, for all *n*=10 random networks optimized to perform *t*=10 named 2-input target functions, the Pearson correlation coefficient between the target function and the optimized responses. The top row corresponds to the responses shown in (D). The results described in the text define success using a threshold Pearson correlation of 0.85, although the results hold for other threshold values (Figure S6E). For the violin plots, the gray violins show the kernel density estimate of the data distributions, red lines show the median values, and only a random subset of the data is displayed as scattered points.

The versatility of individual dimers in random networks was broadly distributed, with some dimers exhibiting much greater versatility than others (Figure 6B). However, some dimers were capable of fitting over 70% of the one-input target functions solely by adjusting their accessory monomer expression levels (Figure 6B, Figure S6B, Figure S6C). Versatility increased with network size, appearing to saturate at about *m*=6 monomers. In contrast, network connectivity did not appear to impact network versatility (Figure S6A). These results suggest that even random networks can potentially achieve versatile computation.

This analysis focused on the versatility of a single output dimer within a network. However, a feature of natural dimerization networks is that multiple dimers can be biochemically active outputs; for example, different transcription factor dimers often bind to distinct DNA binding sites, activating different sets of downstream genes.^2,91^ We thus reasoned that the ability to use different output dimers in different contexts could further extend the versatility of random dimerization networks.

To test this hypothesis, we re-analyzed the optimization results above, allowing each random network to use different output dimers for each target function. With this additional flexibility, individual random networks were remarkably versatile. Nearly all networks with as few as 8 monomers were able to perform 80% of the functions observed across all 8-monomer networks, and the best networks achieved over 95% of such functions (Figure 6C, Figure S6B, Figure S6C). This expanded form of versatility increased rapidly with network size (Figure 6C) but did not depend on network connectivity (Figure S6A). When switching between two target functions, the abundance of at least one monomer in the network almost always changed by at least 100-fold, but the magnitude of this change was not correlated with the Euclidean distance between the two targets (Figure S6D). This suggests that such versatility does not necessarily confer fragility to small fluctuations in protein expression levels.

Finally, we pushed the limits of our optimization pipeline to test whether random networks could perform two-input target functions solely by optimizing the expression levels of their accessory proteins. Because larger networks require more computational resources to both simulate and optimize, we focused on *n*=10 networks of *m*=20 monomers and measured their ability to perform 10 of the two-input functions previously shown in Figure 4C, including seven two-input logic gates, a ratiometric function, an equality function, and a two-input bump function. The best network tested could perform nearly all 10 target functions (Figure 6D), and all 10 random networks could each perform between 6 and 9 target functions (Figure 6E, Figure S6E), using the Pearson correlation coefficient between the target functions and optimized responses as the metric of success (see Methods). All 10 networks were able to perform the ratiometric, NOR, OR, NIMPLY, and AND functions (Figure 6E, Figure S6F). While other functions, such as XOR and the two-input bump function, appeared more difficult to achieve, all functions were achieved by at least 2 of the 10 tested networks (Figure 6E, Figure S6F). Evidently, even random dimerization networks can perform a broad range of computations.

## Discussion

Across many biological contexts, protein dimerization networks interpret combinations of signals to control differentiation, proliferation, and stress responses. Here, we identify several powerful features of such networks that may explain their prominence in nature. More specifically, we demonstrate that competitive dimerization networks are computationally *expressive*, computing a wide variety of input-output functions, and *versatile*, performing different functions solely by tuning the expression levels of network monomers. Reusing a core set of signaling pathways in this way could enable cell-type-specific signaling in organisms with hundreds of cell types, in which it would be infeasible for each cell type to express unique signaling proteins.^92^ Further, dimerization networks can use multiple monomers as inputs, allowing them to make complex decisions that consider multiple sources of information, as has been observed in many natural systems.^15,74,93,94^ Finally, we found that even networks with random interactions, such as those produced by evolutionary duplication and divergence,^95^ can perform near-complete repertoires of network computations simply by tuning their protein expression levels. Overall, these results could explain the ubiquity of transcription factor dimerization networks in natural signaling pathways.

Many natural networks appear to have sufficient size and connectivity to exhibit high expressivity and versatility. Although one- and two-input expressivity and versatility saturates at a network size of 6 monomers, more difficult tasks require larger networks; for example, 10-monomer networks can compute all three-input logic gates (Figure 5C) and 20-monomer networks can compute two-input functions with random interactions (Figure 6D). By comparison, there are 57 human bZIP transcription factors forming dimerization networks of at least 21 monomers,^4^ and 30-50 bZIP proteins are co-expressed in various cell types (Figure S7). Similar statistics characterize the mouse and *Arabidopsis thaliana* bZIP families, the mouse and human nuclear receptor families, and the *Arabidopsis thaliana* MADS-box family (Figure S7B, Figure S7C). In the context of the results presented here, it thus appears that many natural networks have the potential to exhibit complex, cell-type-specific computations.

Our results suggest specific experiments to understand computation by natural networks. For example, in the *Arabidopsis* low-energy stress response,^44^ bZIP1 and bZIP53 “inputs” are directly upregulated by salt stress in roots or by extended darkness in leaves, while bZIP10 and bZIP25 “accessories” are not.^48,56,96^ However, bZIP10 and bZIP25 still play a critical role in network function; whereas double null *bzip1*/*bzip53* mutants can weakly activate downstream target genes in response to stress, quadruple null *bzip1*/*bzip53/bzip10*/*bzip25* mutants cannot.^48,56^ Our results demonstrate that even knowledge of all protein-protein interaction affinities^43^ is not sufficient to predict how dimerizing transcription factors respond to various inputs, as different accessory monomer abundances can produce vastly different input-output responses (Figure 3, Figure 6). bZIP protein (rather than mRNA) abundances in different *Arabidopsis* tissues have not been quantitatively measured^55^ but could be combined with our model to provide testable predictions for how different expression levels of bZIP10 and bZIP25, such as in roots versus leaves,^43,55^ would affect the relationship between the inputs and output gene expression. Ultimately, this combination of experiments and modeling would provide a predictive understanding of how the bZIP proteins sense and coordinate responses to stress, potentially facilitating the engineering of drought-resistant crops^50,97^ or studies of how this network has evolved over time.^4^

More broadly, fully understanding computation in natural dimerization networks will require the ability to measure the complete distribution of dimers and how it responds to perturbations. Techniques based on proximity labeling^98^ or split-pool barcoding,^99^ for instance, have the potential to reveal the abundances of many different endogenous dimers in plant or mammalian cells in high throughput. Such measurements would disentangle computations performed within the dimer network from those performed by other levels of regulation.

Why are competitive dimerization networks such effective computational systems? Even a single dimerization reaction exhibits nonlinear input-output behavior, which can be accentuated by molecular titration^100,101^ and by chaining multiple dimerization reactions together in “paths.” Maslov and Ispolatov have demonstrated that certain conditions can allow input perturbations to propagate along paths as long as ∼4 dimerization steps.^23^ In more complex networks, many such paths intersect, allowing for complex, non-monotonic responses. For example, in the simple network computing a bump function (Figure 2C), one path favors output dimer formation at medium input concentrations, but another path then disfavors the same dimer’s formation at high input concentrations. In this view, computational versatility is possible because one may tune the accessory monomer expression levels to leverage different paths in a network, thereby achieving different input-output functions.

Not all functions can be computed by dimerization networks. Input signals necessarily decay as they propagate because changing the abundance of an input monomer by *N* molecules can, at most, change the abundance of another dimer by *N* molecules. While enzymatic catalysis or gene transcription could potentially be used to amplify the outputs of network computations, signal decay still poses limits on the complexity of their computations. For example, the inverted bump function (Figure 2D) requires paths of only three monomers, whereas more complex functions, such as the “up-down-up” function shown in Figure S1, require paths of up to four monomers and exhibit smaller output dynamic ranges. Some functions are likely too complex to be performed with a dynamic range larger than the intrinsic noise of the system. Nevertheless, as evidenced by the functions shown here, a wide variety of complex computations can still be computed without reaching this limit. Further, in large networks, a single input might coherently regulate multiple monomers to mitigate the challenge of signal decay. For instance, in *Arabidopsis*, sucrose translationally represses all five S1-family bZIPs.^51,52^

Dimerization networks are versatile: tuning their accessory monomer expression levels modulates their input-input computations (Figure 3, Figure 6). However, the same property could also make these computations overly sensitive to protein expression noise. Our results suggest that networks can balance these opposing properties through a separation of concentration scales. For instance, the input-output computation shown in Figure S5E is robust to modest (<3-fold) perturbations of accessory expression levels, but qualitatively sensitive to larger (>10-fold) perturbations (Figure 3B). More broadly, we found that the expression level of at least one monomer typically changed by at least 100-fold to achieve versatility (Figure S6D). This further suggests that dimerization networks can be robust to small fluctuations in accessory expression levels but versatile upon large changes in accessory expression levels.

Beyond studies of natural networks, our results could be applied to synthetic biology and therapeutic development. Synthetic dimerization networks could be engineered by fusing synthetic transcription factors^102,103^ to dimerization domains, benefiting from “failed” attempts to engineer orthogonal dimerization domains.^104–106^ Such synthetic networks could sense multi-input features of cell state or, in CAR T-cells, detect combinations of multiple cell surface antigens.^107^ Separately, our modeling framework could enable more predictable treatment of networks dysregulated in disease. For instance, non-canonical dimerization of nuclear receptors results in unwanted side effects when treating inflammatory disorders with ligands for glucocorticoid receptor.^108,109^ Understanding the full dimerization network could identify combinations of receptor agonists and antagonists, or even double-headed ligands inducing specific heterodimers,^109^ to treat nuclear receptor disorders while minimally perturbing other dimers in the network.

It can be tempting to regard the complexity of protein interaction networks as an accidental byproduct of duplication and divergence during evolution. However, the field of neural network computation has shown that simple but nonlinear elements, when connected in a complex network, can act as powerful computational systems.^110–112^ Dimerization networks are prevalent across biological signaling pathways and, as seen here, offer powerful computational capabilities. These observations strongly suggest that many-to-many dimerization networks could be used as adaptable, multi-input computers whose specific functions can be readily tailored to diverse cellular needs. This system-level viewpoint, complemented with predictive mathematical models, should facilitate the control of natural cellular functions as well as the engineering of synthetic ones.

### Limitations of this work

In the parameter screens, a wide range of affinity constants was chosen so as to capture many diverse network behaviors (see Methods). However, it is unlikely that real protein networks could exhibit such a wide range of interaction affinities. Thus, we performed a subsequent parameter screen using a restricted range of affinity values (four orders of magnitude) and still observed many complex input-output functions (containing up to 2 local extrema; Figure S3F). Secondly, this work analyzed network behaviors at equilibrium, whereas some natural networks function with fluctuating inputs or components.^113^ However, dynamical simulations suggest that our quasi-equilibrium assumption holds as long as there is a sufficient separation-of-timescales between the dimerization equilibration (minutes to hours) and fluctuations in network components (Figure S5B). Finally, dimerization networks likely possess additional capabilities beyond those examined here. Transcription factor dimers may regulate the expression of network monomers, generating feedback, which could produce dynamic behaviors such as oscillations and multistability.^102,113^ This property could also allow the output of one network to activate the inputs of another. Additionally, multiple dimers could be used as outputs, such as in combination (to compute sums of dimer concentrations) or separately (to compute multi-output functions beyond those analyzed here).

## Acknowledgments

We thank Matthew Langley, Pietro Perona, Jordi Garcia-Ojalvo, Arvind Murugan, Erik Winfree, Lulu Qian, Evan Mun, and James Linton for their conversations and scientific input. We thank Jan Gregrowicz, Martin Tran, Rachael Kuintzle, and Rongrong Du for feedback on the manuscript. This research was supported by the National Institutes of Health R01MH116508, the Allen Discovery Center, a Paul G. Allen Frontiers Group advised program of the Paul G. Allen Family Foundation (award number UWSC10142), the National Institute of Biomedical Imaging and Bioengineering of the National Institutes of Health R01EB030015, and Caltech AI4Science Amazon Web Services cloud computing credits. J.P.G. was supported by the National Science Foundation Graduate Research Fellowship and the Ford Foundation Pre-Doctoral Fellowship. A.S. is grateful for support from a Department of Defense Vannevar Bush Faculty Fellowship. M.B.E. is a Howard Hughes Medical Institute investigator.

## Author Contributions

J.P.G., B.E., and M.B.E. conceived of dimerization network computation. The computational approaches used were designed by all authors and implemented by J.P.G. and M.L. J.P.G. and M.B.E. wrote the manuscript, with input from all authors.

## Declaration of Interests

M.B.E. is a scientific advisory board member or consultant at TeraCyte, Plasmidsaurus, and Spatial Genomics.

## Methods

### Resource Availability

#### Lead contact

Further information and requests for resources should be directed to and will be fulfilled by the lead contact, Michael B. Elowitz.

#### Materials availability

This study did not generate any new reagents.

#### Data and code availability

All simulation data and all original code have been deposited at CaltechDATA and are publicly available as of the date of publication at https://doi.org/10.22002/hxnqz-4gv13. Any additional information required to reanalyze the data reported in this paper is available from the lead contact upon request.

## Method Details

All analysis was performed in Python version 3.8.13. Parameter screens and optimization trials were performed using Amazon Web Services (AWS) c5.4xlarge and c6a.48xlarge EC2 instances, respectively.

### Network simulations

The input-output functions of dimerization networks were simulated using the Equilibrium Toolkit (EQTK, version 0.1.3), a Python-based, computationally efficient numerical solver for systems of reversible biochemical reactions.^114^ To more precisely specify the problem of simulating a network’s input-output function, we consider the vector ***c***, the concentrations of all species in the network, as described more thoroughly in the EQTK documentation (https://eqtk.github.io/user_guide/core_concepts.html). Thus,

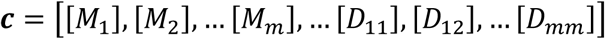

This work presents results in unitless concentrations, as the results would remain the same for any scaling *k* of the concentrations so long as the affinities are also scaled by 1/*k*. Each dimerization network specifies a set of *n*_*dimer*_ chemical reactions in which two monomers dimerize, e.g., *M*_1_ + *M*_1_ ⇌ *D*_11_. All such dimerization reactions can be written as a stoichiometric matrix ***N***, whose rows correspond to dimerization reactions and whose columns correspond to chemical species. Each row specifies how the counts of each chemical species increase or decrease with each chemical reaction.

EQTK seeks to identify the unique set of species concentrations ***c***_*eq*_ at equilibrium. To do this, it imposes two constraints. Firstly, for each reaction involving monomers *i* and *j*, we define the equilibrium (or affinity) constant that must be satisfied:

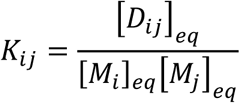

Secondly, we impose the conservation of mass according to the stoichiometry matrix ***N***; the total abundance of all monomers must remain constant. The total concentration of each species is the sum of its free form and all of its dimers:

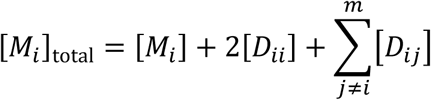

To do this, EQTK defines the conservation matrix ***A*** (using notation consistent with the EQTK documentation; not to be confused with the accessory protein expression levels ***a***), such that the quantity ***A*** ⋅ ***c*** is conserved. For this to be true, ***A*** must satisfy ***A*** ⋅ ***N***^⊤^ = ***0***.

Given the total monomer concentrations as the initial condition, EQTK then uses trust region optimization to identify the equilibrium concentrations of all species consistent with both the dimerization affinities *K*_*ij*_ and the conservation of ***A*** ⋅ ***c***. Thus, to simulate the input-output computation performed by a particular network, the equilibrium species concentrations were solved over a titration of the input monomer(s), holding the total abundance of the accessory (non-input) monomers constant.

### Parameter screen

To perform the large parameter screen, networks of two to twelve monomers were generated, with equal numbers of networks of each possible connectivity. In this parameter screen, approximately 10^6^ networks were generated for one-input simulations, 250,000 of which were used for two-input simulations. For example, a network of eight proteins can have between 7 and 28 heterodimer “edges,” (22 options); thus, for each number of edges, 45,455 networks were generated for a total of 1,000,010 networks. We later sub-sampled these networks to assess whether the number of networks sampled impacted the analysis. When more networks were sampled, the log of the number of unique functions discovered increased linearly with the log of the number of networks sampled (Figure S3C, Figure S3D), but the expressivity trends presented in Figure 4B remained consistent. The networkx Python package (version 2.7.1) was used to randomly generate graphs with a desired number of edges from an Erdős–Rényi model; each graph was checked for connectedness (i.e., that there are no fully separate networks) and re-generated if necessary to achieve connectedness. Homodimer edges were chosen with a probability of 75% (i.e., approximately 75% of monomers across the parameter screen were allowed to homodimerize). This fraction is within the range observed in natural dimerization networks, such as bZIP proteins (70-80%)^4^ and nuclear receptor proteins (68%).^1^ Upon choosing which edges would be present in a network, the values of the edge affinities were randomly chosen on a log-uniform range of dimensionless values 10^-5^ to 10^7^ using Latin hypercube sampling (LHS). Finally, the expression levels of network proteins were also randomly chosen on a log-uniform range of dimensionless values 10^-3^ to 10^3^ using LHS.

The aforementioned parameter ranges were defined generously so as to encompass as many biologically feasible behaviors as possible, including those that may require extreme parameters. However, the parameter ranges are biologically inspired. Protein expression levels in whole mammalian cells, as well as of transcription factors in the nuclei, have been observed to vary over six orders of magnitude,^115–117^ ranging from approximately 5.5 × 10^-13^ M (one copy per cell) to 5.5 × 10^-7^ M (10^6^ copies per cell), assuming cell volumes of approximately 3 pL.^116,118^ The wide range for dimerization affinities was chosen such that the strongest affinity sampled (10^7^) would be strong enough to dimerize 99% of monomers at the lowest concentration sampled (10^-3^), and the weakest affinity sampled (10^-5^) would be weak enough such that only 1% of monomers at the highest concentration sampled (10^3^) would be dimerized. Biological affinity values span an enormous range, with dissociation constants (K_D_) from approximately 10^-3^ M (mM) to 10^-12^ M (pM),^119^ and certain RNase inhibitor proteins have been reported with even stronger affinities (K_D_ < 2 × 10^-16^ M).^120^ While we acknowledge that the affinities characterizing competitive dimerization networks are unlikely to take on such a wide range due to biochemical constraints on affinity and multi-specificity,^121^ many-to-many interactions of SYNZIP coiled-coil proteins have been observed to vary in affinity (K_D_) over four orders of magnitude from approximately 10^-10^ M (100 pM) to 10^-6^ M (1 μM).^104^

All generated parameter sets were simulated over a titration of the input monomer(s), with 30 titration points for one-input functions and 12 titration points for two-input functions; titration points were spaced evenly in log space from 10^-3^ to 10^3^ (the same range as for the accessory monomers). The ray Python package (version 1.11.1) was used for parallelization. For each network, the concentrations of each dimer over this input titration constituted the “responses.” Any concentrations below 10^-3^ were rounded to 10^-3^, as we consider such concentrations outside of the biochemically feasible window (i.e., less than one molecule of dimer per cell). The dataset was filtered to include only responses with a dynamic range greater than 10-fold; all other dimers either did not form at all or did not change significantly in response to the input monomer(s). The remaining responses were categorized into “unique” functions by discretizing the space of possible outputs into ten-fold bins (i.e., bin edges at 10^-3^, 10^-2^, 10^-1^, 1, 10^1^, 10^2^, and 10^3^, for both input and output). For each input bin (e.g., input between 10^-3^ and 10^-2^), the response points were averaged in log space, and this value was categorized into one of the output bins (e.g., 0.5 is categorized into the 10^-1^ to 1 bin). Thus, each response was transformed into a sequence, such as [0, 0, 0, 1, 1, 1], constituting its unique response function. To analyze expressivity, all unique response functions for a given dataset were counted. Lastly, we used these unique functions to create a library of target functions for optimization (see sections below). For each unique function observed, all responses categorized as performing that function were averaged in log space to create the corresponding target function in the library.

To count the number of two-output functions observed in the parameter screen, we iterated through all combinations of dimers in each network, identified which discretized function they were categorized as (using the same discretization scheme as described above), and tabulated the number of unique combinations of discretized functions observed. We compared our results to a scrambled control, in which we randomly sampled the same number of response functions with replacement from our overall dataset, paired them together randomly, and counted the number of unique discretized functions observed. We sampled with replacement because the number of response combinations that needed to be sampled was always larger than the number of responses.

### Dual Annealing Optimizations

Two classes of optimization algorithms were used in this work: (1) a dual annealing algorithm was used to identify optimal parameter sets for particular functions, testing the minimum number of monomers required to achieve each; (2) a genetic algorithm was used in the much larger effort characterizing the versatility of networks with random interaction affinities. In both cases, the ray Python package (version 1.11.1) was used for parallelization.

Dual annealing optimization was used to determine whether networks of a particular size could achieve pre-defined target functions (in Figure 4C) or logic gates (in Figure 5). A dual annealing algorithm was used to optimize the affinities (***K***) and accessory monomer expression levels (***a***) of networks with a defined size to best fit a target function, with the loss defined as the sum of squares of residuals in log space:

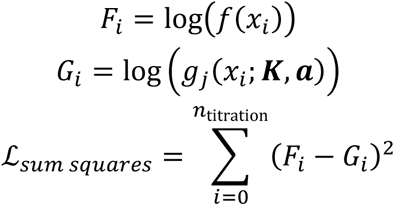

where *F*_*i*_ is the log-scaled target function and *G*_*i*_ is the log-scaled equilibrium concentration *g*_*j*_ of the *j*’th dimer at a particular input concentration *x*_*i*_. Networks are simulated with *n*_titration_ titration points. The optimize.dual_annealing function in the scipy Python package (version 1.10.1) was used to perform such optimizations. The dual annealing algorithm combines simulated annealing with a local search algorithm.^76^ In simulated annealing, a perturbation of the parameter set (a “step”) is proposed based on a visiting distribution. If the proposed step improves the loss function, it is accepted; otherwise, it may still be accepted with some probability based on a “temperature” factor (which decreases over the course of the optimization) so as to promote the identification of globally optimal solutions. In dual annealing, a local search is subsequently performed on the solutions identified by simulated annealing.

To optimize networks for three- and four-input logic gates, four titration points in each input dimension were used, at concentrations of 10^-3^, 10^-1^, 10^1^, and 10^3^. The lower two concentrations were considered “off,” and the higher two concentrations were considered “on.” The optimization was considered successful if the highest response of any input combination in which the output should be “off” is less than the lowest response of any input combination in which the output should be “on.”

### Genetic Algorithm Optimizations

Separately, a genetic algorithm was used to measure the versatility of networks with random binding affinities ***K*** (Figure 6). In this process, we define a library of target functions and evaluate the ability of each sampled network to reproduce each target by optimizing the accessory protein expression levels ***a***. In a genetic algorithm, an initial population of parameter sets is generated and the best of these sets are allowed to “reproduce,” producing a new population of parameter sets. This new population is then “mutated” (perturbed) and the process is repeated. All optimizations in this section were performed using the genetic algorithm (GA) function of the pymoo Python package (version 0.5.0)^122^; all 1-d input functions were optimized using 20 iterations and a population size of 100, and all 2-d input functions were optimized using 200 iterations and a population size of 1000.

For each network size *m*, we defined a library of *N*_*m*_ target functions *f* from the unique functions that were identified in the parameter screen (see “Parameter screen”). Note that each network size *m* thus has a distinct library of size; this allows us to study versatility as a fraction of functions we know to be possible for a given network size.

We consider three loss metrics for evaluating the ability of a network, with affinities ***K*** and accessory monomer expression levels ***a***, to perform a target *f* using dimer index *j*:

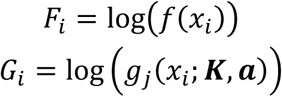

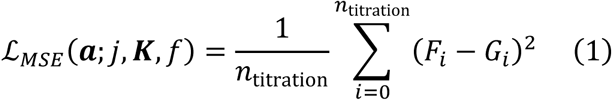

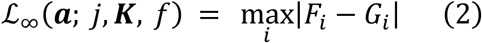

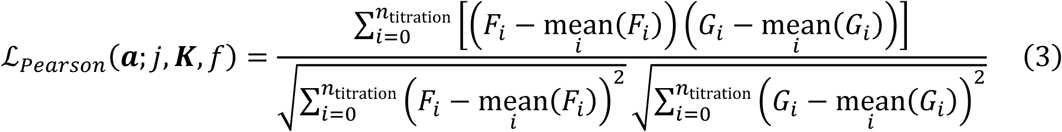

where *F*_*i*_ is the log-scaled target function and *G*_*i*_ is the log-scaled equilibrium concentration *g*_*j*_ of the *j*’th dimer at a particular input concentration *x*_*i*_. Networks are simulated with *n*_titration_ titration points.

In our experiments, we tune ***a*** in order to optimize the mean squared error ℒ_*MSE*_ (Equation 1) but evaluate the quality of the resulting fit using the infinity norm ℒ_∞_ (Equation 2, for one-input functions) or ℒ_*Pearson*_(Equation 3, for two-input functions). This was done because ℒ_∞_(also known as the Chebyshev distance) measures the loss at the “worst point,” which is the strictest metric for whether a response would be acceptable in practice. That is, we have the following:

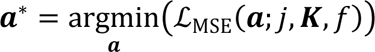

We then define a *versatility* metric *V*_*m*_ measuring the fraction of targets each network could perform with a loss below the tolerance γ = 1. For a given ***K*** ∈ *R*^*d*^, where *d* is the number of dimers, and each particular *j*’th dimer, versatility is calculated as

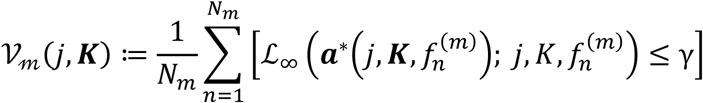

We term *V*_*m*_ as versatility because it reports the fraction of target functions that dimer *j* in a network with affinities ***K*** can perform simply by tuning ***a***. This definition requires that the same dimer must always be used, such that no “rewiring” would be required at the molecular level.

We sampled 50 networks from ***K*** ∼ *U*_*log*_([10^−7^, 10^5^]^*d*^) and subsequently performed the necessary inner optimizations of *a* to measure *V*_*m*_ for each dimer.

In Figure 6C, we broaden our definition of versatility to allow a network to perform different target functions using different output dimers. In this case, we first redefine versatility as the following:

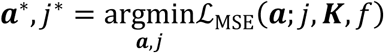

which yields a versatility metric *V* that is independent of dimer index:

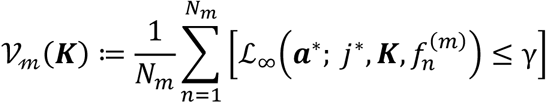

This metric allows us to quantify the ability of a single network ***K*** to perform different functions by tuning ***a*** when given the freedom to use different dimers for each function.

When measuring the versatility of two-input functions, we found the ℒ_∞_ loss function to be unnecessarily strict, as even a one-pixel shift in the response function could increase the loss beyond the threshold γ. As such, for two-input functions, we used ℒ_*Pearson*_ to compare the target function to simulated responses in a more holistic manner. We tested a variety of other metrics as well; we found that the structural similarity index measure (SSIM) and the Wasserstein distance gave results similar to the Pearson correlation, whereas the Hausdorff distance and ℒ_*MSE*_ loss, like the ℒ_∞_ loss, were unnecessarily strict. For the results described in the text, a threshold Pearson correlation of 0.85 was used to assess whether a target function was achieved, although our results hold for different choices of this threshold (Figure S6E).

### Simulating the Kinetics of Network Re-equilibration

Dimerization network re-equilibration kinetics were simulated by numerical integration of ordinary differential equations (ODEs) using the integrate.odeint function of the scipy Python package (version 1.10.1). The ODEs were of the following form:

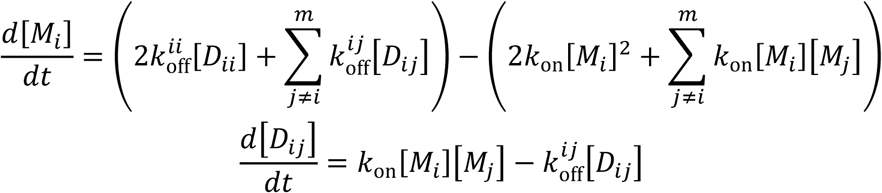

The rate of change in the concentration of monomer *M*_*i*_ is the rate of all dimer dissociation reactions involving monomer *M*_*i*_ minus the rate of all dimer association reactions involving monomer *M*_*i*_, adjusted for stoichiometry. The rate of change in the concentration of dimer *D*_*ij*_ is the rate of the *D*_*ij*_ association reaction minus the rate of the *D*_*ij*_ dissociation reaction. We assume that all association reactions have similar rate constants, following a minimal kinetic model in which there is a single high-energy transition state for dimerization. In this model, various dimers only differ in the free energy of their dimerized states and thus only differ in their dissociation rates. We chose an association rate constant of 5 × 10^5^ M^-1^ s^-1^, following experimental measurements of coiled-coil dimerization kinetics.^123^ The dissociation rate constants were chosen randomly on a log scale between 10^-4^ and 1 s^-1^, giving dimerization affinity constants (K_D_) of about 200 pM to 2 μM, matching experimental measurements of coiled-coil interaction affinities.^104^ These real-unit K_D_ values correspond to ***K*** values in our dimensionless units ranging from about 10^-3^ to 10^1^. All parameters were made unitless by converting concentration units into “molecules per cell” counts, assuming a cellular volume of about 3 pL.^116,118^

20 random networks were simulated for each network size from *m*=2 to *m*=12 monomers. The initial state of each network was the equilibrium state in which the input monomer was at its lowest concentration (1 molecule per cell), but the concentration of the input monomer was then set to its maximum concentration (10^6^ molecules per cell). Networks were simulated with time increments of 10 s until equilibrium, with a maximum simulation time of 10^7^ s. The exact equilibrium concentrations were calculated using EQTK (see “Network Simulations”), and a network species was defined as having reached equilibrium when its concentration was within 1 molecule per cell of the exact equilibrium concentration.

To assess whether the dimerization reactions can be at quasi-equilibrium despite changing input abundances, we simulated network dynamics as the total concentration of one monomer was oscillated sinusoidally. The oscillating monomer was made to have a total concentration oscillating between 1 and 10^6^ molecules/cell, using the same affinity parameters and concentration conventions as above. *n*=10 different networks of each network size from 2 to 12 monomers were simulated. For each dimer, we compared the dynamical trajectory of its concentration to the calculated equilibrium concentration of the dimer at each timepoint. We calculated the fraction of dimers for which the dynamical and equilibrium concentrations matched at every timepoint within 0.5 log units (i.e., with less than a ∼3-fold difference).

### Assessing the Intrinsic Noise of Dimerization Equilibria

The intrinsic noise of the dimerization equilibria was simulated using stochastic Gillespie simulations, implemented using a modified version of the code from the biocircuits python package (version 0.1.14), using the numba Python package (version 0.55.1) to speed up simulation functions and the ray Python package (version 1.11.1) for parallelization. In the Gillespie algorithm, a population of molecules as well as a set of possible reactions among those molecules (with rates calculated from the current population) is first defined. To perform a step of the simulation, the time until the next reaction is sampled from an exponential distribution based on the rates of all reactions. The identity of the reaction that occurs is chosen based on the relative rates of all the possible reactions, and the population is updated to reflect that reaction.

Using the same parameter ranges as in the deterministic simulations of network equilibration, 20 random networks were simulated for 1000 s for each network size from *m*=2 to *m*=12 monomers. The noise η in the concentration of each species was defined as the coefficient of variation in the number of molecules per cell *n* over time^80,82^:

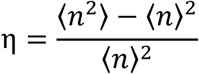

### Assessing Robustness to Extrinsic Monomer Expression Noise

Protein expression levels are subject to both intrinsic expression noise, which independently affects different genes, as well as extrinsic expression noise, in which overall fluctuations in machinery for transcription and translation could affect the expression of all genes in a concerted manner. The steady-state distribution of mRNA counts produced by bursty transcription is negative binomial,^124^ and steady-state protein concentrations appear similarly distributed.^82^ Thus, expression noise was modeled here using a gamma distribution, the continuous analog of the negative binomial distribution, using the probability density function below:

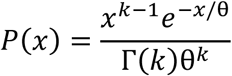

where *k* is the shape parameter, θ is the scale parameter, and Γ(*k*) is the Gamma function of *k*. This distribution was parameterized to produce expression noise coefficients similar to those observed experimentally (0.4 for intrinsic and 0.6 for extrinsic).^82^ To accomplish this, the shape parameter was set to 1/η^2^ and the scale parameter was set to η^2^.

We simulated both cases in which there was either purely intrinsic (independent) or purely extrinsic (concerted) expression noise. For each unique function identified in the aforementioned parameter screen, a network performing that function was randomly selected and simulated for 50 random perturbations (independent or concerted, for intrinsic or extrinsic noise) of the accessory monomer expression levels. Each of the 50 resulting input-output functions (simulated at 30 input points) was compared to the original input-output response using the root-mean-square difference (RMSD) in log space:

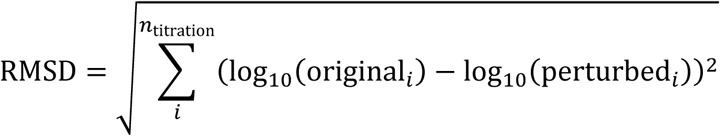

The RMSD is one metric for the typical difference, in log space, between the original and perturbed responses at each input point.

### Transcriptomics data for analysis of transcription factor co-expression

To characterize how many dimerizing transcription factors are potentially co-expressed in individual cell types, two pre-existing datasets were used: the integrated mouse transcriptomics dataset from Granados et al.,^21^ which integrated multiple pre-existing mouse datasets, and the Human Protein Atlas (version 23.0) “RNA consensus tissue gene data” dataset^125^ (https://www.proteinatlas.org), which reports normalized expression levels from a consensus of multiple scRNA-seq datasets. The Human Protein Atlas dataset was used to demonstrate the cell-type-specific expression of nuclear receptor proteins in Figure 1C. The names of genes belonging to the bZIP and nuclear receptor families in mice and humans were obtained from Uniprot,^126^ Reinke et al.,^4^ and Amoutzias et al.^1^

### Quantification and Statistical Analysis

The numbers of samples (*n*) used in each analysis are described in the Method Details section as well as the figure captions. In analyses involving the comparison of data distributions, such as the comparisons of versatility scores in Figure 6, violin plots were used to show the distribution of data points. More specifically, violins show the kernel density estimate calculated using the kdeplot function of seaborn (version 0.12.2). Points were also directly plotted with the violins; points were sub-sampled if it was not practical to display all data points. On such violin plots, the medians of the data are displayed as red lines.

## Additional Resources

An interactive notebook hosted on Google Colaboratory, in which users can simulate the input-output functions of arbitrary dimerization networks, can be found at the following link: https://bxky.short.gy/interactive_dimerization_networks

## Key Resources Table

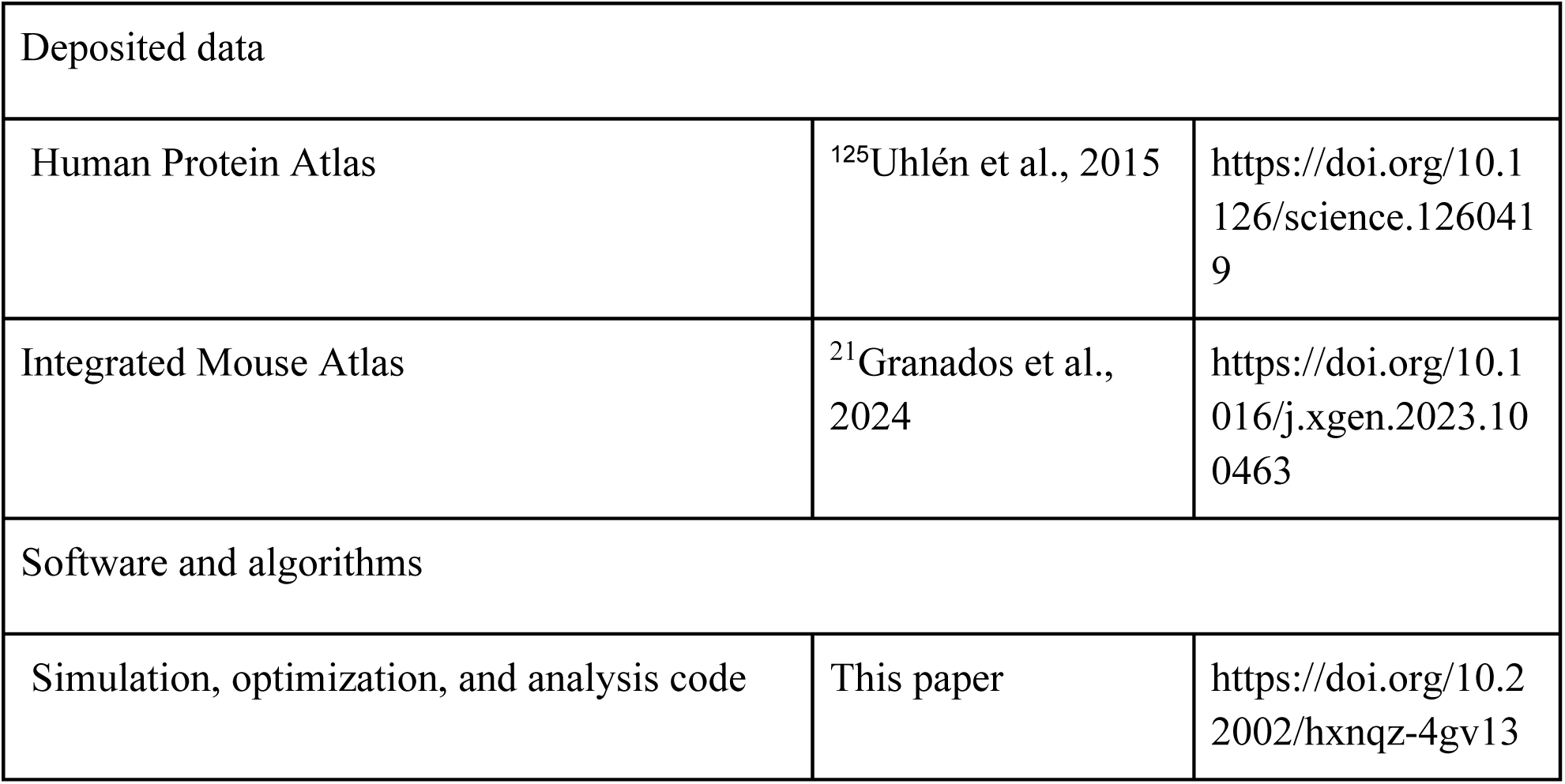

## Supplemental Figures

**Figure S1.**
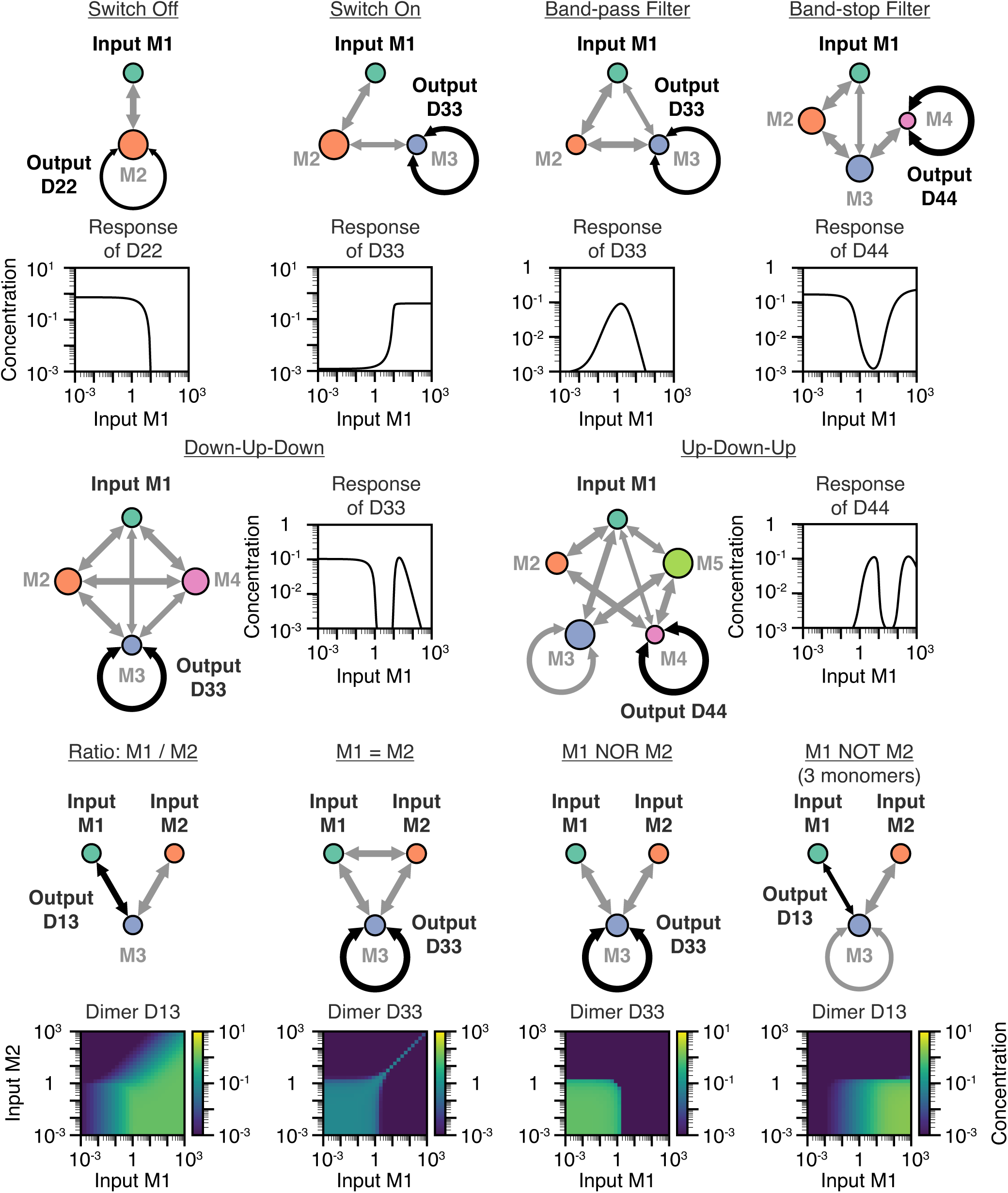
An atlas of elementary dimerization network computations, part 1. Shown for each function is a schematic of the network parameters and a plot of the corresponding input-output function, for both one-input functions (top) and a partial set of two-input functions (bottom). The remaining two-input functions can be found in Figure S2. For all panels, the networks shown were inspired by networks from the random parameter screen (Figure 4) and rationally pruned to identify minimal topologies capable of computing each input-output function. All results are displayed in unitless concentrations (see Methods).

**Figure S2.**
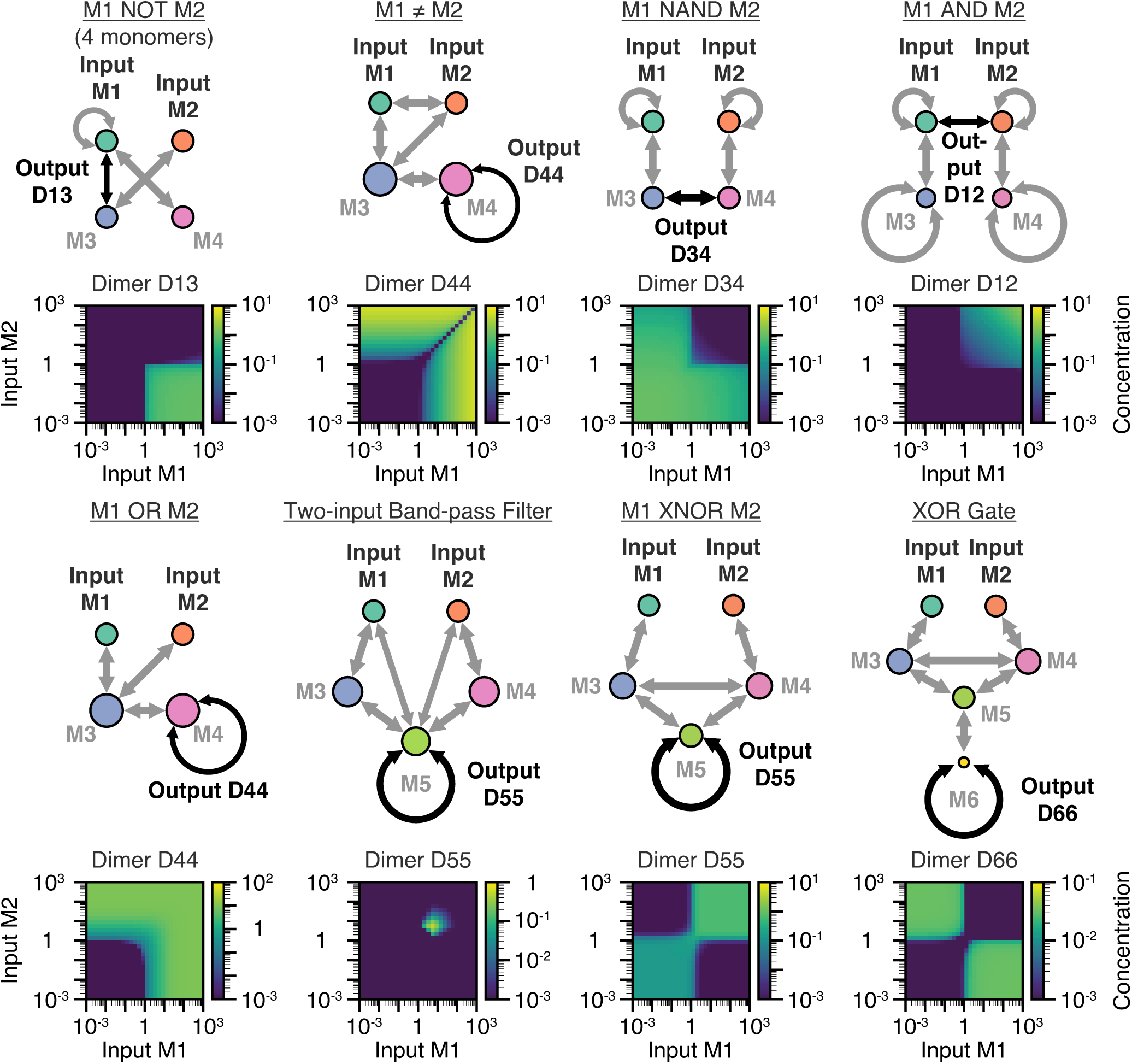
An atlas of elementary network computations, part 2. Shown for each function is a schematic of the network parameters and a plot of the corresponding input-output function. Displayed are the two-input computations not included in Figure S1. For all panels, the networks shown were inspired by networks from the random parameter screen (Figure 4) and rationally pruned to identify minimal topologies capable of computing each input-output function. All results are displayed in unitless concentrations (see Methods). These computations can be rationally understood. For instance, considering the XOR network, heterodimerization of M3 and M4 in the absence of either input allows M5 to heterodimerize with M6, limiting the formation of the D66 output dimer. Adding either input alone sequesters either M3 or M4, allowing the other of the two to dimerize with M5, freeing M6 to form the D66 output dimer. When both inputs are present, though, sequestration of both M3 and M4 allows M5 to again dimerize with M6, reducing the D66 output.

**Figure S3, related to Figure 4.**
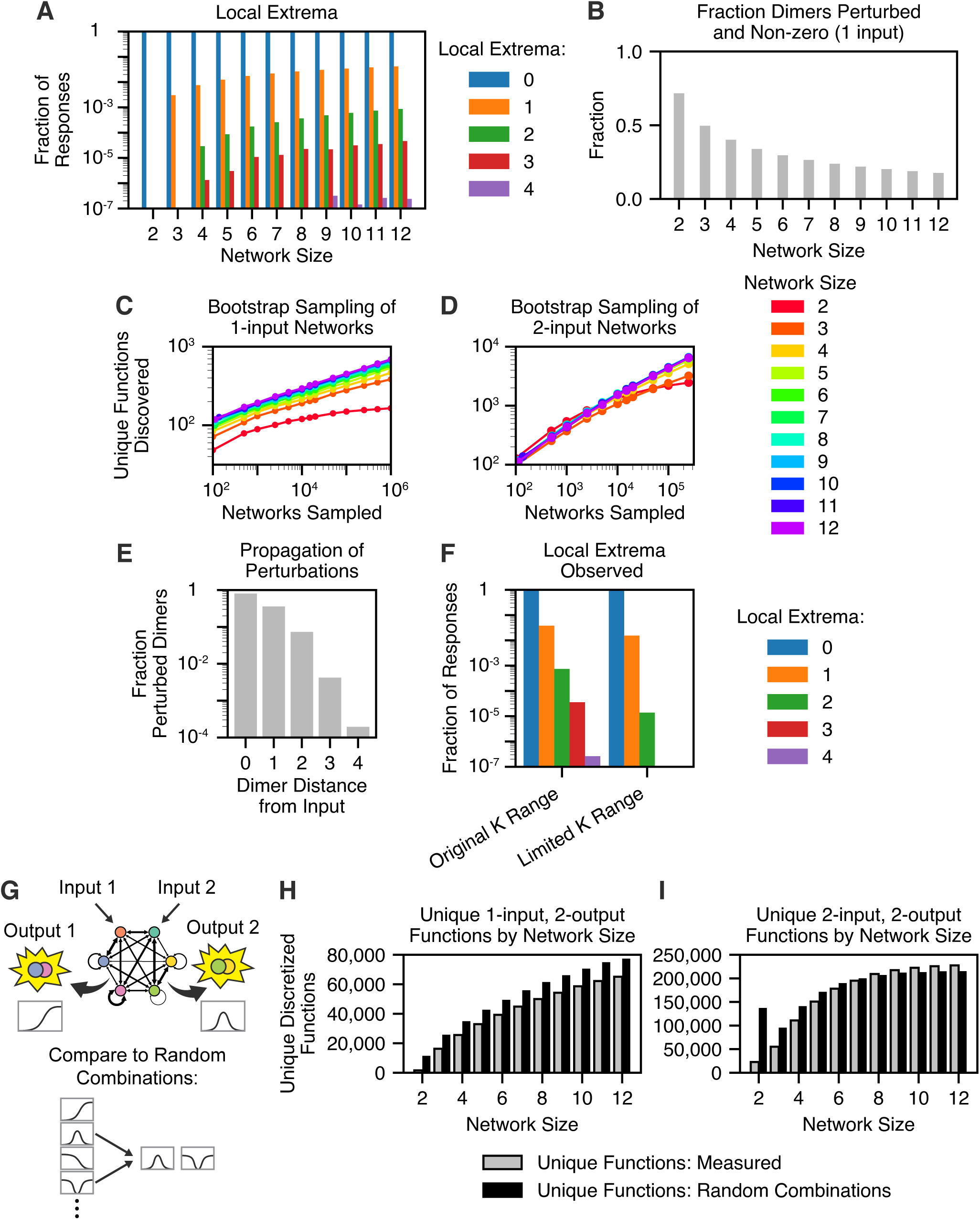
A global parameter screen reveals the diversity and nature of dimerization network computations. (A) A bar graph shows, for each network size, the fraction of responses with zero to four local extrema (i.e., local minima and maxima). (B) A bar graph shows, for each network size, the fraction of dimers that both form at significant concentrations and are perturbed more than 10-fold by a titration of the input monomer. (C-D) The number of unique one-input (C) and two-input (D) functions observed is plotted versus the number of networks sampled in the random parameter screen. (E) A bar graph shows, for increasing distances between the input monomer and output dimer, the fraction of dimers (out of all dimers that form at appreciable concentrations) that are perturbed more than 10-fold in response to a titration of the input monomer. (F) A bar graph shows, for a parameter screen of 12-monomer networks using a more limited range of affinities (K_ij_) from 10^-3^ to 10^1^, the fraction of responses with zero to four local extrema (i.e., local minima and maxima). (G) A schematic depicting how two dimers within the same network could be used to compute two-output functions. (H-I) Bar graphs show, for each network size, the number of unique, discretized, one-input, two-output functions (H) or two-input, two-output functions (I), as well as the number of unique functions for a scrambled control in which random pairs of response functions were selected from the overall dataset. The outlier for the two-input *m*=2 scrambled data appears to be due to the *m*=2 dataset having a more even distribution of unique functions among the whole set of responses.

**Figure S4, related to Figure 5.**
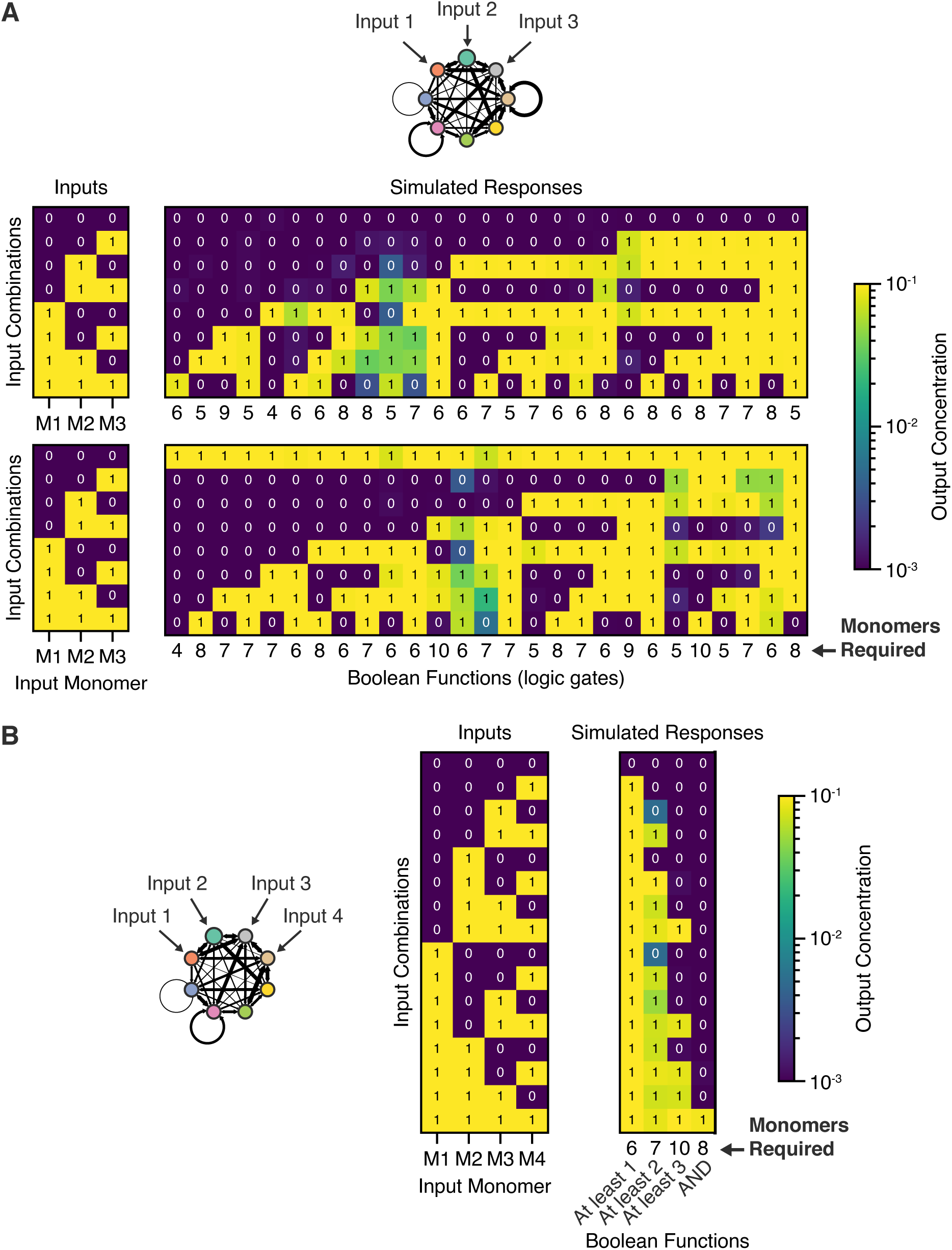
Competitive dimerization networks can compute multi-input functions. (A) Dimerization networks can perform all three-input logic gates. (left) Rows represent different combinations of inputs that are presented to each network. (right) A heatmap of responses is shown, where each column represents a unique logic gate and the color of the response heatmap represents the output dimer concentration. The number of network monomers required to perform each gate is noted below each column. (B) Dimerization networks can further perform four-input logic gates. Shown are four examples, the AND and “at least *n*” gates, which output 1 if at least *n* inputs are present.

**Figure S5.**
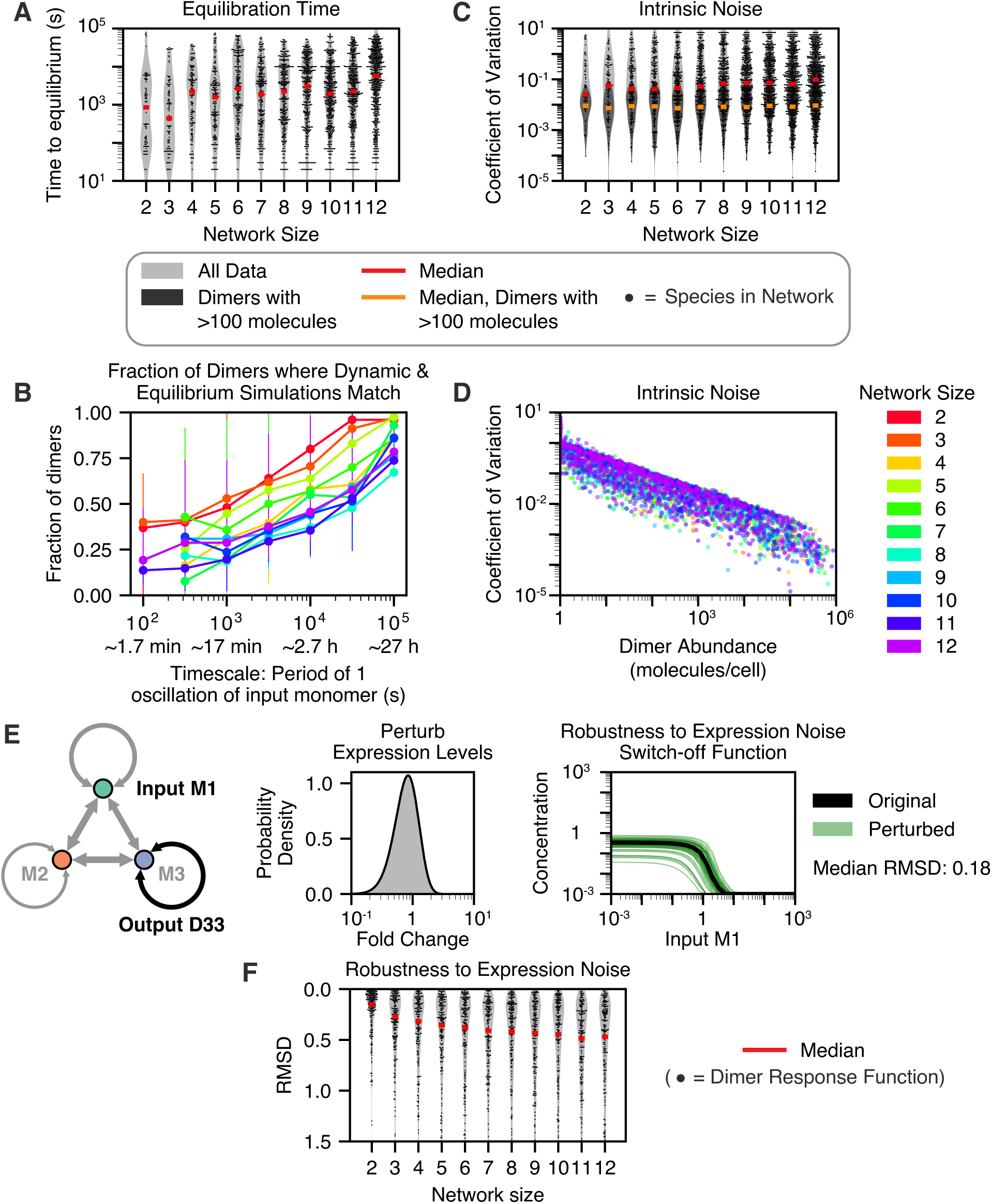
Competitive dimerization networks exhibit biologically reasonable equilibration kinetics and robustness to noise. (A) Network equilibration kinetics were simulated by numerically integrating ordinary differential equations (ODEs) describing dimer association and dissociation kinetics. The time for each species in *n*=20 networks to re-equilibrate after a perturbation of an input monomer is displayed as a violin plot with scattered points. (B) To assess the timescale at which a dynamical dimerization network can no longer be assumed to be at equilibrium, we simulated network equilibration as the total concentration of one monomer was oscillated sinusoidally. For each dimer, we compared the dynamical trajectory of its concentration to the calculated equilibrium concentration of the dimer at that timepoint. Shown is the fraction of dimers (out of all dimers whose concentrations change significantly over the course of the simulation) for which the dynamical and equilibrium concentrations matched at every timepoint (within 0.5 log units, or less than ∼3-fold difference), for various network sizes *m* as well as different timescales at which one monomer was perturbed sinusoidally. *n*=10 different networks of each network size were simulated; the error bars show the 1^st^ and 3^rd^ quartiles of the data across different networks. Points for the 100 s timescale with network sizes 4-10 were not shown, as 40-70% of these simulations failed numerically. (C-D) The intrinsic noise of the binding equilibrium was simulated using the Gillespie algorithm with 100 steps of 10 s each. (C) A violin plot (light gray) with scattered points shows the coefficient of variation, a measure of noise, for each species. A dark gray violin shows the data specifically for species present at high abundances (>100 molecules/cell, median shown by the orange line). (D) A scatterplot shows the relationship between the equilibrium abundance (in molecules/cell) and the intrinsic noise measured for each simulated species. (E) The versatile switch-off function from Figure 3B (right, black) was perturbed with typical protein expression noise (probability density function shown in the middle), and each perturbed computation is shown (right, green, *n*=50 perturbations). The median root-mean-square deviation (RMSD), in log space, between the original and the perturbed curves was 0.18, corresponding to a 1.5-fold change in output. (F) Networks performing each unique function from the parameter screen were perturbed with noise affecting the expression level of each monomer independently (*n*=50 perturbations). A violin plot with scattered points shows the RMSD in log space between the original and perturbed input-output functions, with an inverted y-axis to emphasize that low RMSD values correspond to high robustness. For all violin plots, the violins show the kernel density estimate of the data distributions, red lines show the median values, and only a random subset of the data is displayed as scattered points.

**Figure S6, related to Figure 6.**
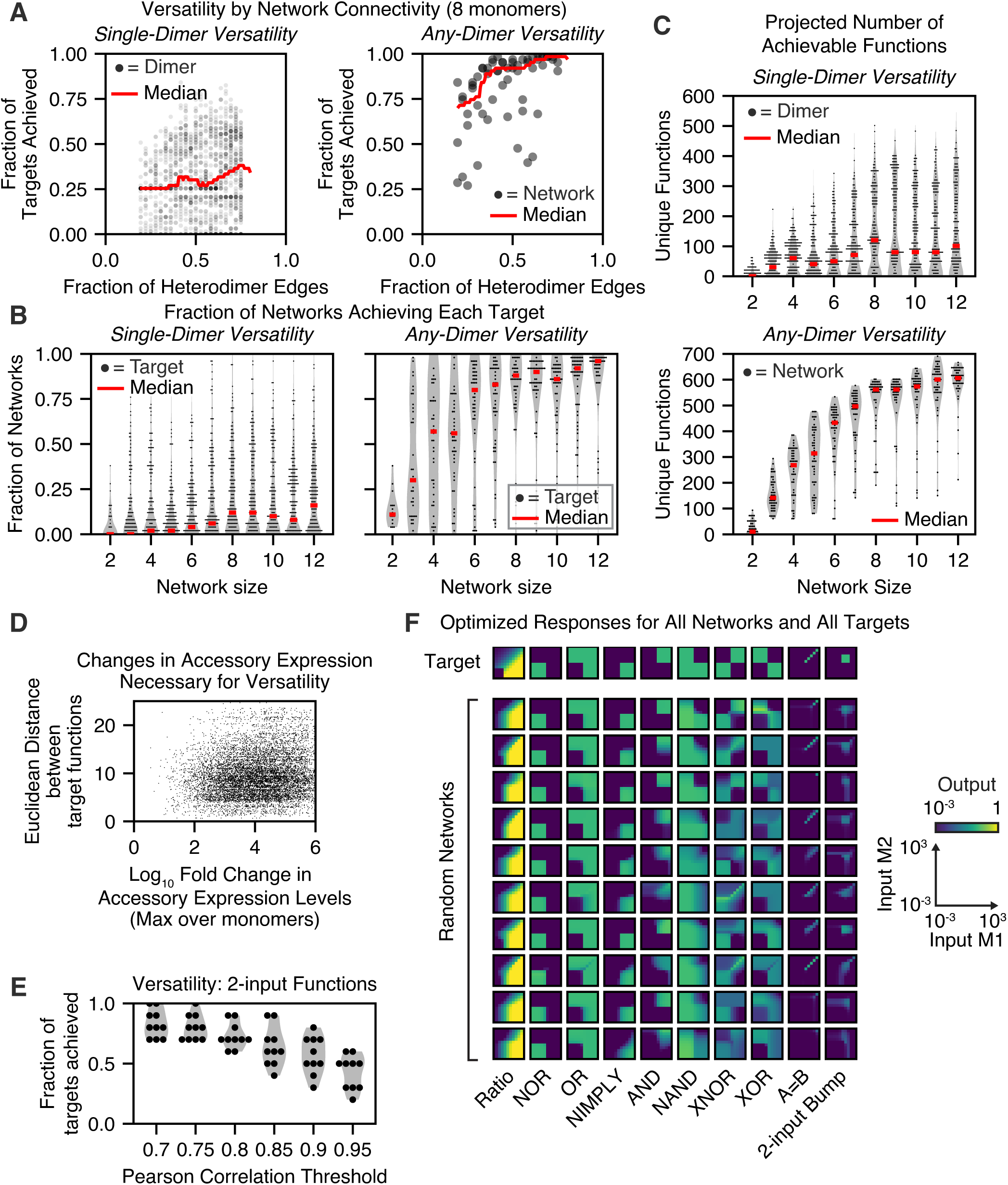
The versatility of random networks increases with network size. (A) A scatter plot shows how network versatility using only a single output dimer (left) or any output dimer (right) varies with network connectivity. (B) To show how difficult each target function was to achieve, a violin plot with scattered points shows the fraction of random networks that could achieve each target function, separated by network size and whether only a single dimer (left) or any dimer (right) may be used as the output. (C) While Figure 6B shows only the fraction of targets that could be achieved by each network, plotted here is the projected total number of targets each network is expected to achieve – accounting for differences in the overall expressivities of networks differing in size – for both cases in which only a single dimer (top) or any dimer (bottom) may be used as the output. (D) A scatterplot showing, for all combinations of target functions achieved in the versatility analysis by networks with *m*=8 monomers, both the maximum log fold change in accessory expression level (maximum over all accessory monomers) and the Euclidean distance between the two target functions. (E) A violin plot with scattered points shows the versatility of *n*=10 random networks with *m*=20 monomers toward *t*=10 2-input target functions at different thresholds of the Pearson correlation coefficient. (F) An array showing the responses of all *n*=10 random networks optimized to perform *t*=10 named 2-input target functions. For the violin plots, the gray violins show the kernel density estimate of the data distributions, red lines show the median values, and only a random subset of the data is displayed as scattered points.

**Figure S7.**
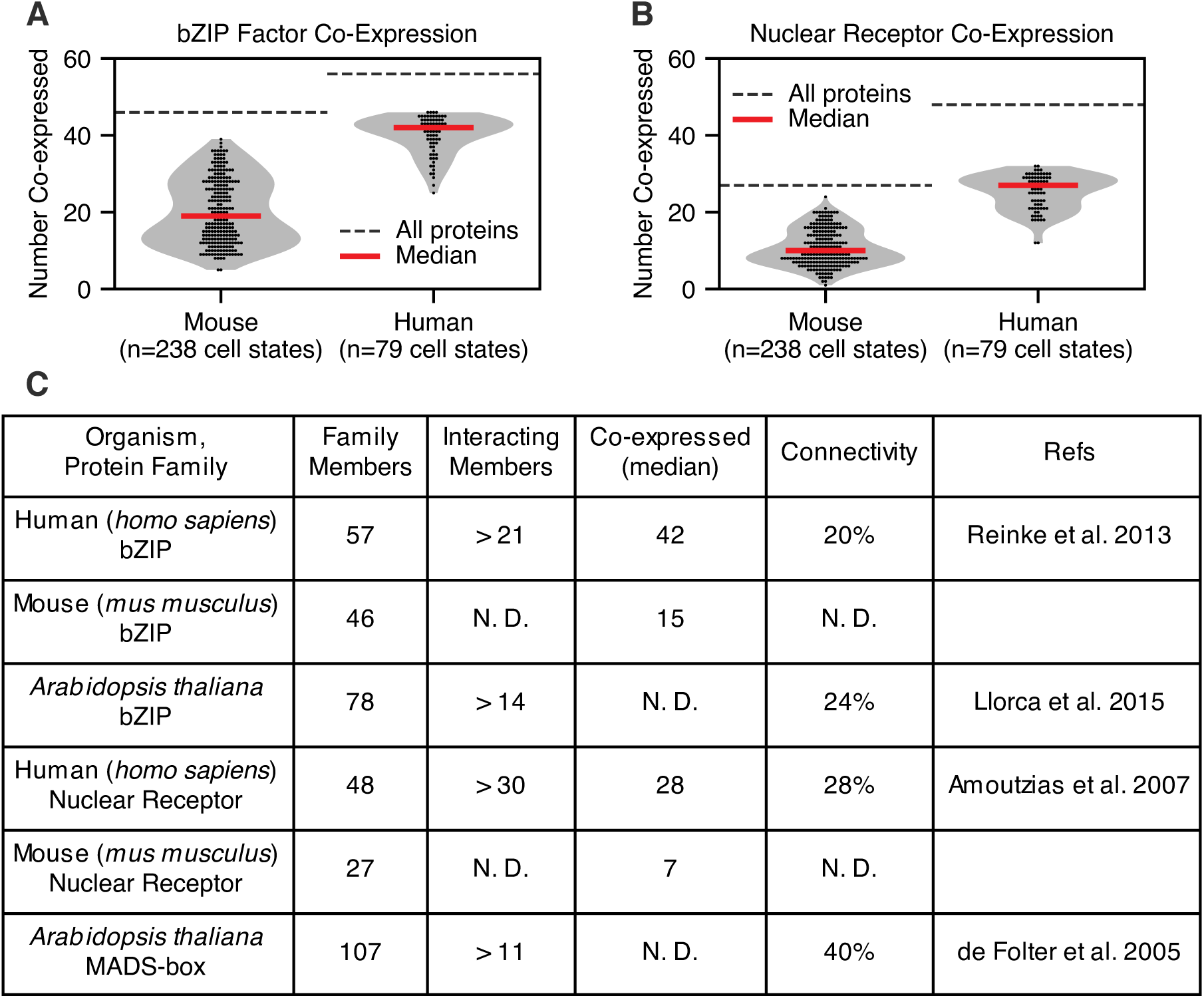
Natural dimerization networks are of sufficient size to exhibit high expressivity and versatility. (A, B) Co-expression of bZIP (A) and nuclear receptor (NR) (B) transcription factors was assessed for many cell types across both mouse and human datasets. A violin plot with scattered points shows the number of network proteins co-expressed in each cell type. Gray violins show the kernel density estimate of the data distributions, red lines show the median values, and black dotted lines indicate the total number of genes assessed. (C) Table summarizing the size, number of known interacting members, number of co-expressed members, and connectivity of several natural dimerization networks. N.D. indicates that an entry was not determined.

Supplemental Video 1: Tunable Two-input Bump function

Supplemental Video 2: Versatile Network for Both AND and NOR Logic Gates

Supplemental Video 3: Versatile Network for Both AND and NAND Logic Gates

## Notes

https://doi.org/10.22002/hxnqz-4gv13

### Summary of Updates

We have added several major computational results to the paper and have added several sections to the text connecting our results to dimerization networks in nature. The new computational results include the following: (1) While our previous results demonstrated that networks with random interaction affinities can be tuned to perform nearly all one-input functions, we have now demonstrated that larger random networks of 20 monomers can also perform a variety of two-input functions, including logic gates such as XOR and XNOR as well as a two-input bump function (Figure 6D, Figure 6E). (2) To assess the extent to which dimerization networks can function at quasi-equilibrium despite dynamically changing inputs, we performed dynamical simulations of network computations with oscillating concentrations of input at varying frequencies. We determined that most dimers are at quasi-equilibrium when the inputs change on physiologically relevant timescales of hours to days (Figure S5B). (3) We measured the expressivity of two-output functions using two separate output dimers. We found that, for both one and two inputs, dimerization networks can perform nearly all possible two-output functions (i.e., nearly all combinations of unique functions observed in the parameter screen) (Figure S3G, Figure S3H, Figure S3I). (4) To assess whether versatile networks can still be robust to small fluctuations in accessory expression levels, we calculated the extent to which accessory expression levels changed for networks to exhibit versatility for one-input, one-output functions. We found that most networks changed the expression of at least one accessory monomer by at least 100-fold to change the identity of the output function (Figure S6D), suggesting that these networks would still be robust to intrinsic noise in accessory protein expression (<3-fold).

